# Innate Immune Remodeling Drives Therapy Resistance via Macrophage–NK Cell Crosstalk

**DOI:** 10.1101/2025.07.22.666055

**Authors:** Chia-Hsin Hsu, Keng-Jung Lee, Jingyi Chen, Leanne R. Donahue, Jianping Lin, Danielle Kacaj, Zhong-Yin Zhang, Andrew C. White

**Author notes:** Corresponding author. Andrew C. White, 618 Tower Rd., T3-014A VRT, Ithaca, NY 14850.

## Abstract

The dynamic evolution of the immune tumor microenvironment (TME) during targeted therapy is a critical yet poorly understood determinant of treatment response and resistance. While most studies compare immune states before and after treatment, temporal immune changes during therapy remain largely uncharacterized, limiting development of effective combination strategies. Here, we investigated immune dynamics throughout targeted therapy using mouse melanoma models that recapitulate human therapeutic responses. Single-cell RNA sequencing (scRNA-seq) identified a previously unrecognized inflection point where the inflamed TME during tumor regression, characterized by robust NK cell infiltration, transitions to an immune-excluded state upon onset of drug-tolerant residual disease. We uncovered a unique macrophage subset (F4/80^hi^CCL5⁺MHCII⁺CD63⁺) that orchestrates NK cell recruitment through CCR2/5 signaling during regression. Depletion of these macrophages using LysM-cre;iDTR mice significantly reduced NK cell infiltration. Specifically during residual disease, pharmacological inhibition of Ptpn22, a phosphatase that negatively regulates immune activation, reprogrammed macrophages, restored NK cell recruitment and enhanced therapeutic efficacy. Extending these findings to human cancer, longitudinal scRNA-seq analysis of melanoma and lung cancer patient samples revealed dynamic NK cell infiltration during targeted therapy, establishing a direct link between innate immune remodeling and treatment outcome. Unlike prior prognostic studies assessing immune states at single time points, our results provide mechanistic evidence of a temporal relationship between NK cell infiltration and therapeutic efficacy. Together, these findings position immune evolution as a driver of acquired resistance and identify macrophage–NK cell crosstalk as a therapeutically actionable axis to overcome immune exclusion and improve targeted therapy across multiple cancer types.

**One Sentence Summary:** Cancer therapy resistance emerges from a dynamic evolution of the tumor microenvironment, characterized by macrophage-driven NK cell infiltration during initial tumor regression, followed by exclusion of NK cells during residual disease, highlighting macrophage-NK cell interactions as a promising therapeutic target to improve clinical outcomes.

## INTRODUCTION

Targeted therapies have shown remarkable efficacy across various cancer types by disrupting aberrant kinase activation. However, drug resistance remains an almost universal challenge. Beyond their effects on tumor cell proliferation and survival, targeted therapies also dynamically alter the immune tumor microenvironment (TME), enhancing antigen presentation, MHC expression, and infiltration of CD4+ and CD8+ T cells while reducing regulatory T cells (Tregs) *(1–15)*. These enhanced T cell-mediated effects have laid the groundwork for combining targeted therapies with immune checkpoint blockade (ICB), such as anti-PD-1/PD-L1 therapies *(2, 10–13, 16, 17)*. Unfortunately, such combinations often lead to increased toxicity without substantially improving patient outcomes, highlighting the urgent need for alternative strategies to enhance therapeutic efficacy *(18–22)*.

While previous studies have explored TME alterations induced by targeted therapies, they have largely focused on comparisons between pre-treatment and post-treatment tumors or between treatment-sensitive and resistant tumors. Whether these therapy-induced TME alterations persist throughout treatment, whether the immune-excluded TME emerges before resistance develops, or whether the TME undergoes dynamic changes during therapy remains unclear. Addressing this gap is critical for understanding how targeted therapies influence tumor evolution and resistance development.

An immune-inflamed TME, characterized by robust infiltration of immune cells, particularly cytotoxic CD8+ T cells, is associated with active anti-tumor responses and favorable outcomes *(23, 24)*. While adaptive immunity plays a vital role in therapeutic efficacy, accumulating evidence suggests the need to engage a coordinated immune response involving both innate and adaptive systems *(25–27)*. Innate immunity often serves as a critical initiator of T cell responses by providing signals that drive their activation and recruitment *(28, 29)*. Moreover, powerful methods of immune-mediated cancer cell destruction exist that do not depend on T cells, but rather on innate immune cells *(30–32)*. An in-depth understanding of the TME’s cellular composition, dynamics, and functional state during therapy is essential for uncovering novel therapeutic targets and overcoming resistance mechanisms.

To address these gaps, we utilized melanoma models that exhibit profound regression upon MAPK pathway inhibition (BRAF/MEK inhibitors, BRAF/MEKi), followed by resistance development with continued treatment. These models closely recapitulate therapeutic responses observed in human cancers. Here, we reveal the temporal evolution of the TME during therapy, demonstrating a transition from an immune-inflamed to an immune-excluded phenotype as tumors acquire resistance, highlighting the importance of maintaining or restoring an immune-inflamed TME to improve therapeutic efficacy. We identify NK cell infiltration as a defining feature of the immune-inflamed TME and a critical determinant of therapeutic efficacy. Mechanistically, we show that NK cell recruitment is driven by CCR2/5 signaling, which originates from a distinct macrophage population characterized by elevated MHC II expression and CCL chemokine secretion. Furthermore, we demonstrate a proof-of-concept therapeutic strategy for manipulating NK cells and macrophages to reinstate anti-tumor immunity. Finally, we validate our pre-clinical findings by confirming a positive correlation between NK cell infiltration and therapeutic responses in human melanoma samples treated with BRAF/MEKi and in human lung cancer patients receiving tyrosinase kinase inhibitors (TKIs).

## RESULTS

### The therapy-induced immune-inflamed TME is positively correlated with targeted therapy efficacy

Analysis of publicly available transcriptomic data from paired biopsies of melanoma patients, taken before and during MAPK inhibition treatment *(33)* revealed increased expression of the pan-leukocyte marker CD45 (PTPRC), suggesting immune modulation in response to targeted therapies (Fig. 1A). To evaluate whether the immune system is essential for targeted therapy efficacy, we employed the YUMM1.7 mouse melanoma model (*Braf^V600E^, Pten^-/-^Cdkn2a^-/-^*) in NSG (NOD-SCID IL2Rγ^null^) mice treated with BRAF/MEKi. In immunodeficient NSG mice, BRAF/MEKi treatment temporarily suppressed tumor growth but failed to induce significant regression, and tumors rapidly regrew within two weeks (Fig. 1B). In contrast, the same model in immunocompetent mice recapitulated key features of human responses to targeted therapy, including tumor regression, establishment of drug-tolerant residual disease, and tumor re-growth due to acquired resistance after 3 – 4 weeks (Fig. 1C and 1D). These results indicate that while targeted therapies are designed to inhibit oncogenic signaling, their full efficacy requires an intact host immune system.

**Figure 1.**
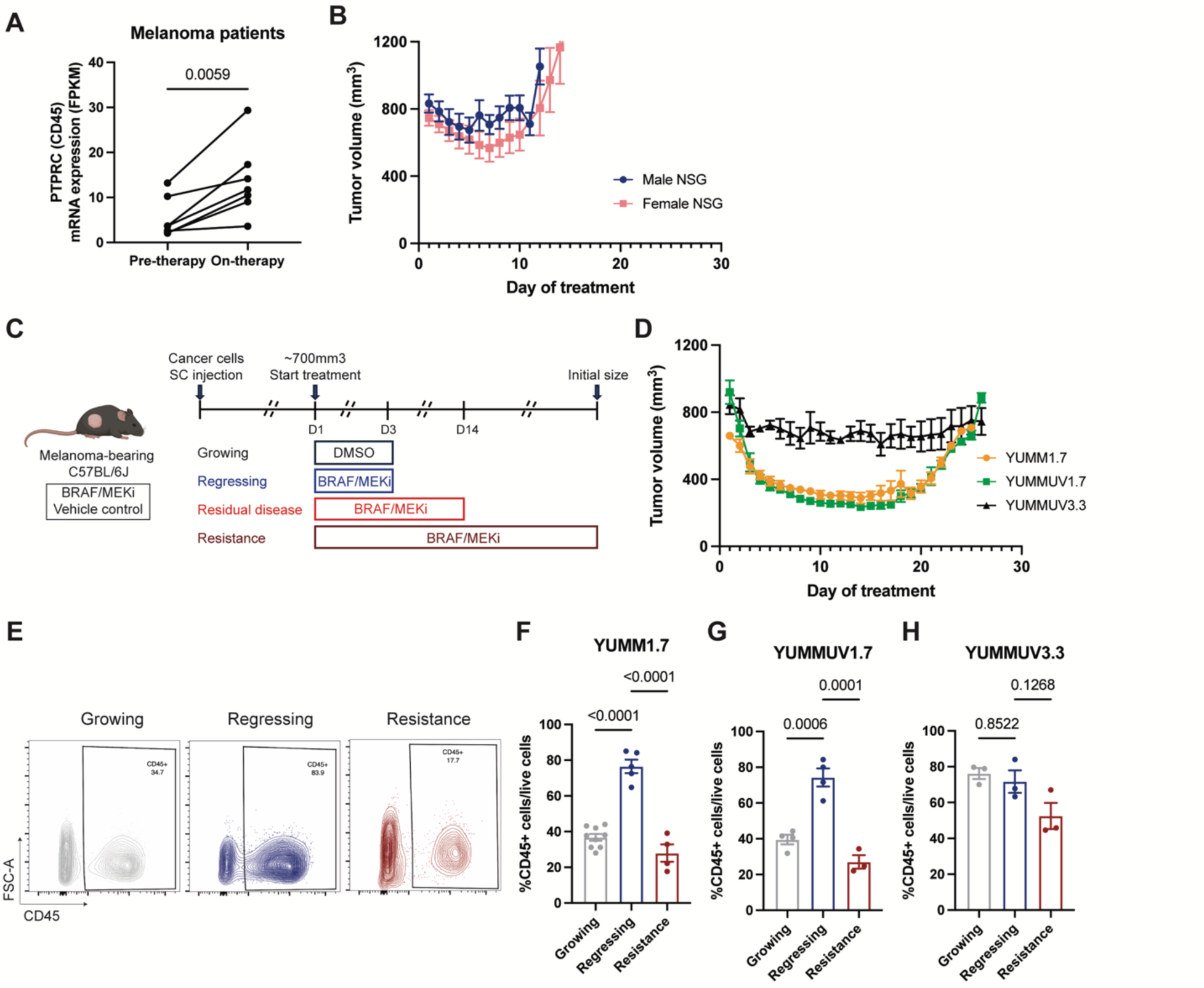
Therapy-induced immune-inflamed TME is positively correlated with targeted therapy efficacy. (A) Dot plots comparing pre-therapy and on-therapy CD45 (PTPRC) mRNA levels in melanoma patients treated with BRAFi or BRAF/MEKi. Matched patient sample size of n = 6. Statistics were calculated using a two-tailed paired t-test. (B) Average tumor volume curves in response to daily BRAF/MEKi in YUMM1.7 melanoma-bearing NSG mice. Tumor volumes were measured daily. (n = 5-6 tumors, mean ± SEM) (C) Schematic illustrating the different phases of syngeneic melanoma C57BL6/J mouse models. BRAF/MEKi was administered by daily oral gavage when tumors reached ∼700mm3 in size. Growing tumor: 3-day vehicle control treatment. Regressing tumor: 3-day BRAF/MEKi treatment. Residual disease: 14-day BRAF/MEKi treatment. Resistant tumor: BRAF/MEKi treatment until the rebounding tumor reaches its initial size. Created with BioRender.com. (D) Average tumor volume curves in response to daily BRAF/MEKi in different melanoma C57BL6/J models. Tumor volumes were measured daily. (n = 3-5 tumors, mean ± SEM) (E) Representative flow cytometry plots of CD45+ cells gated on live cells in different phases in BRAF/MEKi-treated C57BL/6J mice bearing YUMM1.7 (growing tumors were treated with vehicle control). The numbers in the plots represent the percentage of cells within each gate. (F-H) Quantification of CD45+ cell infiltration in different phases in BRAF/MEKi-treated C57BL/6J mice bearing YUMM1.7 (D), YUMMUV1.7 (E), and YUMMUV3.3 (F) (growing tumors were treated with vehicle control). YUMMUV3.3 tumors did not exhibit profound regressing phase as YUMM1.7 and YUMMUV1.7 but the collection timepoint was after 3-day BRAF/MEKi, consistent with timeline defined in (C). (n = 3-9 tumors, one-way ANOVA with Tukey’ s multiple comparisons test, mean ± SEM)

To explore immune dynamics during therapy, we analyzed tumor-infiltrating CD45⁺ immune cells. BRAF/MEKi treatment significantly increased tumor-infiltrating CD45+ immune cells, indicating a therapy-induced immune-inflamed TME. However, this phenotype was lost as tumors acquired resistance (Fig. 1E and 1F). We observed similar dynamics in the YUMMUV1.7 model, an ultraviolet-irradiated version of YUMM1.7, with a higher somatic mutation burden (34) further supporting the association between therapeutic efficacy and immune infiltration (Fig. 1D and 1G). In contrast, the YUMMUV3.3 model (BrafV600E, Cdkn2a–/–) (34) which showed limited tumor regression upon BRAF/MEKi treatment (Fig. 1D), exhibited no therapy-induced immune TME changes compared to untreated tumors (Fig. 1H). Together, these findings suggest that therapy-induced immune-inflamed TMEs synergize with the anti-proliferative effects of targeted therapies to achieve superior tumor control.

### Tumor regression and residual disease are characterized by distinct immune cell signatures

To define the transition from an immune-inflamed to an immune-excluded TME, we sought to precisely deconvolute the immunological landscape in regressing tumors and drug-tolerant residual disease using single-cell RNA-seq (scRNA-seq) on immune cells from tumors (Fig. 1B and 2A). Drug-tolerant residual disease represents a population of persister cells that survive drug exposure while most cancer cells are rapidly eliminated. These persisters are thought to give rise to tumor relapse and regrowth *(35)*. Focusing on this residual disease time point, rather than fully resistant tumors, provides valuable insights into the early mechanisms of drug resistance before the evolution of diversified resistance trajectories typically seen in regrowing tumors. Immune cells were isolated using CD45 microbeads. Enrichment was validated by flow cytometry with a purity of > 95% (Fig. S1A). We analyzed 41,588 single immune cells, with 20,052 and 21,536 immune cells from regressing tumors and residual disease, respectively. Within these profiles, we successfully identified seven immune cell populations based on transcript signatures, including T cells, B cells, NK cells, monocytes/macrophages, dendritic cells (DC), neutrophils, and mast cells (Fig. 2B, 2C and S1B).

**Figure 2.**
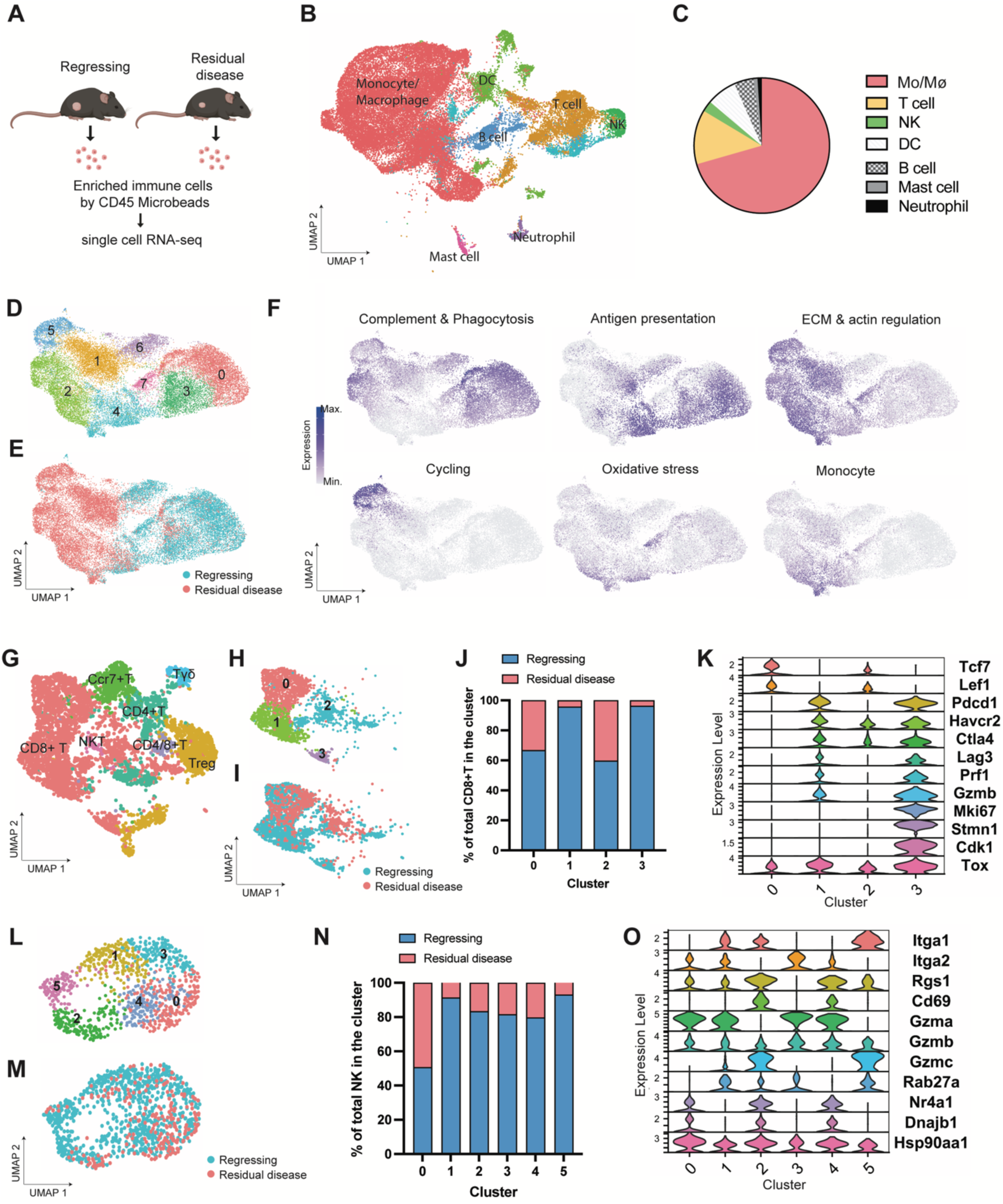
On-therapy BRAF/MEKi-tolerant residual tumors show distinct immune cell signatures compared to on-therapy regressing tumors. (A) Experimental schematic for single-cell RNA-seq. Immune cells from regressing tumors and residual disease were enriched with CD45 microbeads for single-cell RNA-Seq. Created with BioRender.com. (B) UMAP plot of the cell populations obtained from all sequenced CD45+ cells. (C) Pie chart showing the percentage of each immune cell type in (D). Mo/Mϕ, monocyte/macrophage. (D and E) UMAP plots showing re-clustering of the monocyte/macrophage cell type, defining eight clusters (D) from the two analyzed tumor phases (E). (F) Visualization of gene set scores associated with different biological processes in the macrophage clusters. (G) UMAP plot showing annotation of T cell subtypes. (H–J) UMAP plots showing re-clustering of CD8+ T cells defining four clusters (H) from the two analyzed tumor phases (I). Percentage of CD8+ T cells from regressing tumors and residual disease within each cluster (J). (K) Expression of selected genes for stem-like, terminally exhausted effector, precursor-exhausted, and cycling markers in each CD8+ T cell cluster. (L–N) UMAP plots showing re-clustering of NK cells defining six clusters (L) from two analyzed tumor phases (M). Percentage of NK cells from regressing tumors and residual disease within each cluster (N). (O) Expression of selected genes encoding tissue-retained, cytotoxic, and stress response markers in each NK cell cluster.

We were particularly interested in macrophages, NK cells, and CD8+ T cells since macrophages are the most abundant cell type among all the immune cell populations in both regressing tumors and residual disease, and NK and CD8+ T cells are the primary cytotoxic cells against cancers (Fig. S1C and S1D). After re-clustering the sequenced macrophages from both groups, we identified eight distinct clusters based on transcript signatures (Fig. S2A), with macrophages from regressing tumors and from residual disease showing distinct distributions across the clusters (Fig. 2D and 2E). Notably, ∼85% of the macrophages from regressing tumors were distributed in Clusters 0, 3, 6, and 7, compared to only 8% of the macrophages from residual disease (Fig. S2B). Clusters 0, 3, 6, and 7 consisted of macrophages mainly from regressing tumors (Fig. S2C). In contrast, the majority of macrophages from residual disease were distributed within Clusters 1, 2, and 5 (Fig. S2C). Cluster 4 consisted of a similar percentage of cells from both regressing tumors and residual disease (Fig. S2C). Next, we investigated the biological processes between different macrophage subpopulations in regressing tumors and residual disease. Applying gene sets of different macrophage signatures reported by previous studies *(36, 37)*, we observed that Clusters 0 and 3 were enriched for genes associated with complement, phagocytosis, and antigen presentation. Clusters 1, 2, 5 and 7 were enriched for genes involved in ECM and actin regulation, with Clusters 5 and 7 particularly enriched for cell cycle and oxidative stress genes, respectively. Cluster 4 displayed a monocyte signature, including classical and nonclassical, as well as increased MHC class II expression, in line with reported monocyte-to-macrophage transitions (Fig. 2F, S2D-S2F) *(36)*.

To analyze CD8+ T cells, we re-clustered T cells to annotate the T cell subtypes (Fig. 2G and S3A), and found that CD8+ T cells were significantly decreased in residual disease (Fig. S3B). By sub-clustering CD8+ T cells, we identified four CD8+ T cell subpopulations based on transcript signature (Fig. 2H and S3C). All clusters were decreased in residual disease compared to regressing tumors, with Clusters 1 and 3 being significantly reduced (Fig. 2I, 2J and S3D). The CD8+ T cell subpopulations included stem-like (Cluster 0, with co-expression of *Tcf7* and *Lef1*), precursor-exhausted (Cluster 2, with co-expression of *Tcf7*, *Ctla4*, and *Tox* but low expression of *Gzmb*), terminally exhausted effector (Cluster 1, with co-expression of *Gzmb*, *Tox,* and multiple inhibitory receptors, including *Pdcd1*, *Havcr2*, *Ctla4*, and *Lag3*), and cycling CD8+ T cells (Cluster 3, with co-expression of *Mki67*, *Stmn1*, and *Cdk1*) (Fig. 2K) *(38)*.

Similar to CD8+ T cells, NK cells were significantly diminished in residual disease, exhibiting almost four times fewer tumor-infiltrating NK cells than in regressing tumors (Fig. S3E). To identify the difference in NK subtypes in both treatment conditions, we re-clustered the NK cells and identified six subtypes based on transcript signature (Fig. 2L and S3F). Of note, all NK clusters were decreased in residual disease, except for Cluster 0 (Fig. 2M, 2N and S3G). We next investigated the potential functions of each NK subtype. Previous studies have reported several tissue-resident NK cell markers, such as *Rgs1*, *Cd69*, *Itga1* (*Cd49a)*, *Cd103*, and *Cxcr6 (39, 40)*. Clusters 2 and 5 exhibited enriched tissue-resident genes (i.e., *Itga1* and *Rgs1*) and highly expressed *Gzmc,* but not *Gzma* (Fig. 2O). On the other hand, Cluster 3 was the only population with little to no expression of *Itga1*, *Rgs1* and *Cd69,* but with high expression of *Itga2 (CD49b) (41)*, suggesting this cluster to be freshly infiltrating NK cells (Fig. 2O). Tumor-associated NK cells or dysfunctional NK cells have been characterized as exhibiting elevated stress response, reduced cytotoxicity, and higher expression of *Klra* families (equivalent to KIRs in humans), and other genes, such as *Dnajb1* and *Nr4a1(39)*. Markers of dysfunctional NK cells, such as *Klra4/7/8/9*, *Hsp90aa1*, *Dnajb1* and *Nr4a1* were highly expressed in Clusters 0 and 4 (Fig. 2O and S3H). Although expressing *Gzma* and *Gzmb*, these two clusters expressed low levels of *Rab27a*, a key player in NK cell degranulation, suggesting potentially impaired cytotoxicity (Fig. 2O) *(42, 43)*.

Collectively, our scRNA-seq analysis revealed a distinct immune cell atlas between therapy-induced regressing tumors and therapy-tolerant residual disease. Macrophages were the most abundant cell type in both regressing tumors and residual disease, although distinct macrophage populations were observed. Multiple clusters were identified in the CD8+ T and NK cell groups, with a reduction observed in almost every cluster in the context of residual disease.

### NK cells, but not CD8+ T cells or macrophages, are required for optimal targeted therapy response

Since the scRNA-seq data revealed dynamic changes in macrophages, CD8+ T cells, and NK cells during residual disease, we hypothesized that these cell types play significant roles in the tumor response to BRAF/MEKi (Fig. 3A). To identify the immune cell type that contributes to the tumor response to BRAF/MEKi, we evaluated the time required for YUMM1.7-bearing mice to achieve full resistance after depletion of macrophages, CD8+ T cells, or NK cells.

**Figure 3.**
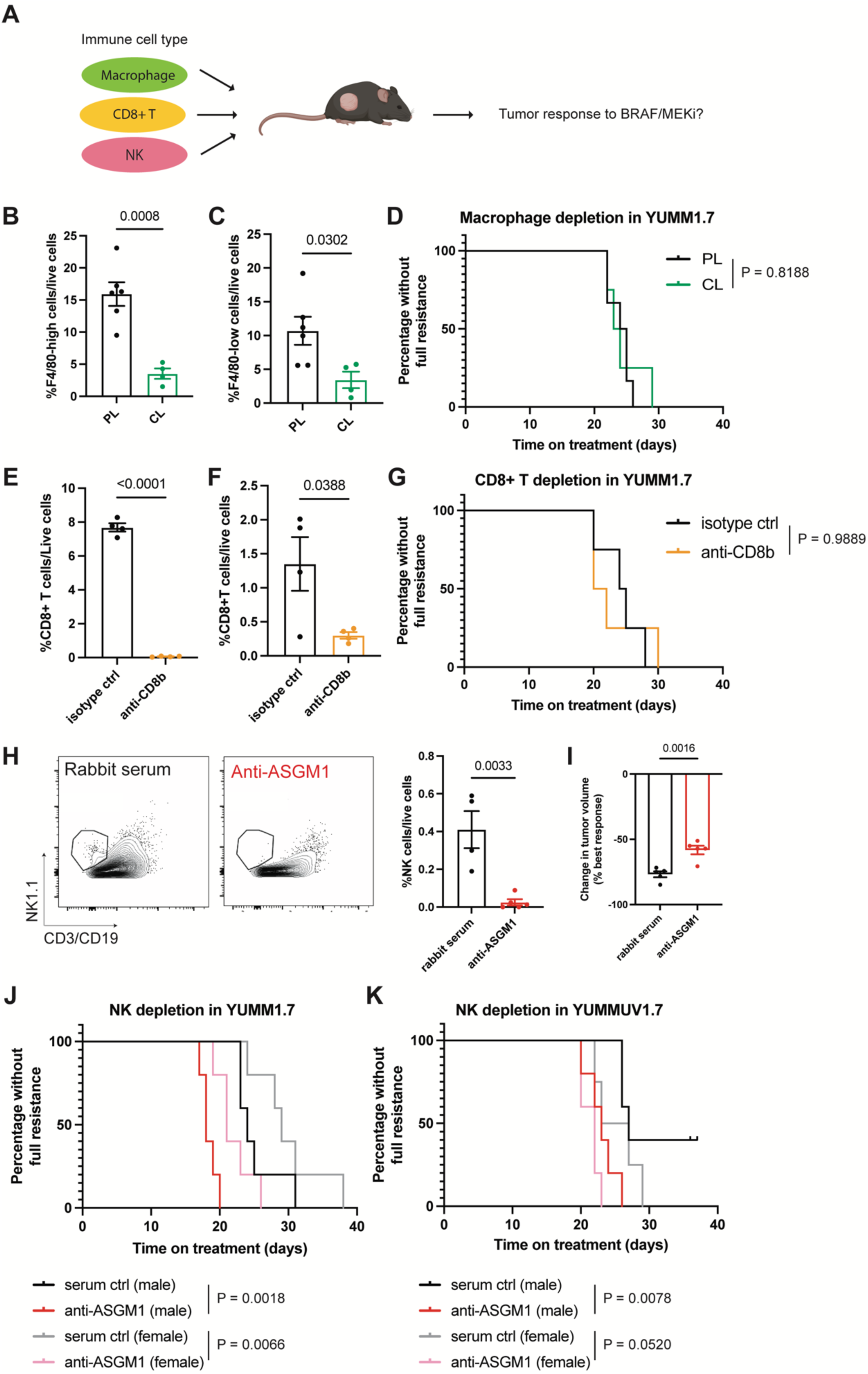
NK cells are essential for BRAF/MEKi therapeutic efficacy. (A) Schematic showing immune cell types, including macrophages, CD8+ T cells, and NK cells, that may contribute to the BRAF/MEKi tumor response. Created with BioRender.com. (B and C) Depletion efficiencies of clodronate liposomes in tumors at the endpoint (full resistance). Quantification of F4/80-high macrophages (B) and F4/80-low macrophages (C). (n = 4-6 tumors, two-tailed unpaired t-test, mean ± SEM) (D) Kaplan-Meier curve for BRAF/MEKi-treated mice bearing YUMM1.7 receiving clodronate liposomes (CL) or PBS liposomes (PL). (n = 4-6 tumors, Log-rank Mantel-Cox test) (E and F) Depletion efficiencies of the anti-CD8b antibody. Quantification of circulating CD8+ T cells six days post anti-CD8b antibody injection (E) and tumor-infiltrating CD8+ T cells at the endpoint (full resistance) (F). (n = 4 tumors, two-tailed unpaired t-test, mean ± SEM) (G) Kaplan-Meier curve for BRAF/MEKi-treated mice bearing YUMM1.7 receiving anti-CD8b antibody or isotype control. (n = 4 tumors, Log-rank Mantel-Cox test) (H) Depletion efficiency of the anti-ASGM1 antibody in tumors at the endpoint (full resistance). Representative flow cytometry plots and quantification of tumor-infiltrating NK cells. (n = 4-5 tumors, two-tailed unpaired t-test, mean ± SEM) (I) Bar plot showing best response (%) of BRAF/MEKi-treated male mice bearing YUMM1.7 receiving the anti-ASGM1 antibody or rabbit serum. (n = 5 tumors, two-tailed unpaired t-test, mean ± SEM) (J and K) Kaplan-Meier curve for BRAF/MEKi-treated male and female mice bearing YUMM1.7 (J) or YUMMUV1.7 (K) tumors receiving the anti-ASGM1 antibody or rabbit serum. (n = 4-5 tumors, Log-rank Mantel-Cox test)

Since macrophages constitute the most prevalent immune cell type and display distinct populations in regressing and residual disease, we postulated that macrophages in regressing tumors and residual disease fulfill disparate roles in resistance onset. We hypothesized that macrophages in residual disease adopt a more pro-tumor-like phenotype. To test this hypothesis, we depleted macrophages using clodronate liposomes starting from day 10, during the establishment of residual disease, while continuing daily BRAF/MEKi treatment (Fig. S4A). Surprisingly, even with significant depletion of tumor-associated macrophages (Fig. 3B, 3C and S4B), there was no significant difference in full resistance onset between the clodronate liposome and PBS liposome controls (Fig. 3D, S4C and S4D).

For CD8+ T cell depletion, an anti-CD8b neutralizing antibody efficiently depleted both circulating and tumor-infiltrating CD8+ T cells (Fig. 3E, 3F, S4E and S4F). However, there was no statistically significant difference in the onset of full resistance between the depletion and isotype control groups (Fig. 3G, S4G and S4H).

For NK cell depletion, an anti-asialo GM1 (ASGM1) antibody was administered intraperitoneally once every four days until full resistance (Fig. S5A). Flow cytometry confirmed the depletion of tumor-infiltrating NK cells (Fig. 3H and S5B). In the presence of anti-ASGM1, tumors more rapidly developed full resistance to BRAF/MEKi. This result was observed in both male and female mice (Fig. 3J, S5C-S5F). Notably, in male mice bearing YUMM1.7 with NK cell depletion, the tumors responded to BRAF/MEKi less effectively, resulting in a larger volume at maximum regression (Fig. 3I). The finding that NK cells strongly contribute to the BRAF/MEKi response was not limited to the YUMM1.7 mouse model. Using YUMMUV1.7, we observed a similar outcome. Tumors in both male and female mice exhibited accelerated development of full resistance when NK cells were depleted (Fig. 3K, S5G-S5J).

### Therapy-tolerant residual disease exhibits reduced NK cell infiltration without altered susceptibility to NK cell cytotoxicity

Our scRNA-seq analysis revealed a significantly decreased quantity of tumor-infiltrating NK cells in residual disease compared to regressing tumors (Fig. 2M and S3E). Furthermore, the NK cell depletion data indicated a more rapid onset of full resistance (Fig. 3J and 3K), suggesting the critical involvement of NK cells in the tumor response to BRAF/MEK inhibitors. To further evaluate the role of NK cells in the control of resistance onset, we examined the level of tumor response to BRAF/MEKi (in % tumor volume change) and the number of tumor-infiltrating NK cells during several treatment phases, including growing tumors (treated with vehicle control), regressing tumors, residual disease, and tumors developing resistance to BRAF/MEKi. We observed a positive correlation between NK cell infiltration and tumor response, with NK cell numbers increasing significantly during the regressing phase, decreasing during residual disease, and remaining low in resistant tumors (Fig. 1C, 4A-4D).

In addition to the number of tumor-infiltrating NK cells, previous studies have shown that NK cells may become exhausted in tumors *(44, 45)*. Thus, we analyzed our scRNA-seq data to determine whether tumor-infiltrating NK cells become exhausted or less activated in residual disease. Our analysis did not reveal any significant differences in the transcript signatures of NK cells between regressing tumors and residual disease (Fig. 4E). There was a similar expression level of various NK activating markers including Granzyme B (*Gzbm*), Cd107a (*Lamp1*), Nkp46 (*Ncr1*), and Perforin (*Prf1*), and exhaustion markers such as Tim3 (*Havcr2*), PD-1 (*Pdcd1*), *Tigit*, and Nkg2a (*Klrc1*), between regressing tumors and residual disease (Fig. 4E-4G). Flow cytometry analysis of these markers corroborated these findings, indicating that there was no significant increase in NK cell exhaustion in residual disease compared with regressing tumors (Fig. S6A-S6D), suggesting that tumor-infiltrating NK cells did not become more exhausted during therapy-tolerant residual disease than during the regressing phase.

**Figure 4.**
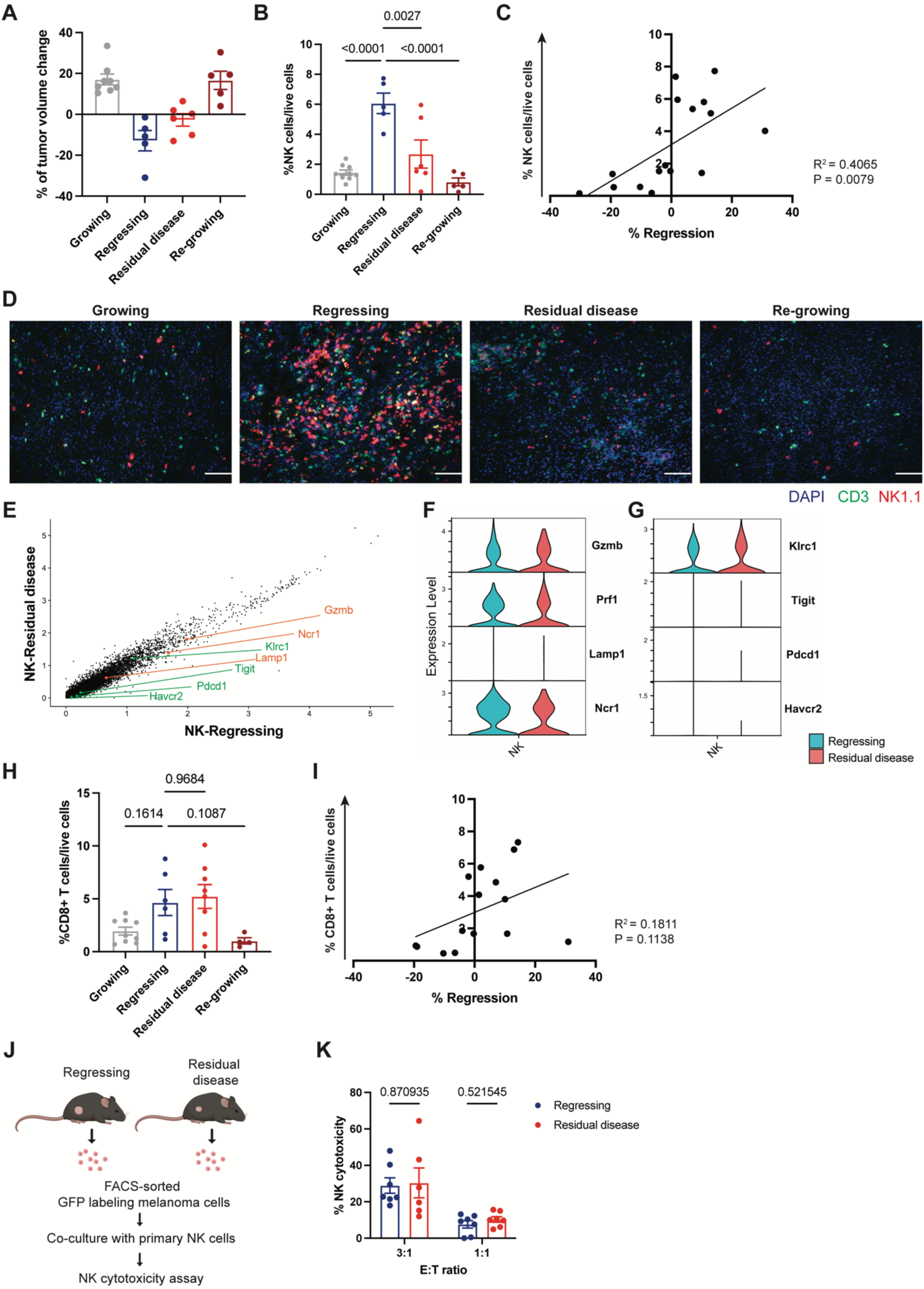
Therapy-tolerant residual disease exhibits reduced NK cell infiltration without altered susceptibility to NK cell cytotoxicity. (A) Bar plot showing tumor volume change (%) of different phases in BRAF/MEKi-treated mice bearing YUMM1.7 (growing tumors were treated with vehicle control). (n = 5-8 tumors, mean ± SEM) (B) Quantification of NK cell infiltration in different phases in BRAF/MEKi-treated mice bearing YUMM1.7 (growing tumors were treated with vehicle control). (n = 4-9 tumors, one-way ANOVA with Tukey’s multiple comparisons test, mean ± SEM) (C) Correlation between NK cell infiltrate in YUMM1.7 tumors and therapy response to BRAF/MEKi. (Simple linear regression) (D) Representative immunofluorescence images of NK1.1 (red), CD3 (green) and nuclear DAPI (blue) of BRAF/MEKi-treated tumors in the indicated treatment groups. NK cells are NK1.1+CD3-. Scale bar 100 μm. (E) Scatter plot showing the NK cell signature between regressing tumors and residual disease. NK cell activation and inhibitory marker genes are shown in orange and green, respectively. (F and G) Violin plot showing the expression of activation (F) and inhibitory (G) marker genes in NK cells from regressing tumors and residual disease. (H) Quantification of CD8+ T cell infiltration in different phases in BRAF/MEKi-treated mice bearing YUMM1.7 (growing tumors were treated with vehicle control). (n = 4-9 tumors, one-way ANOVA with Tukey’s multiple comparisons test, mean ± SEM) (I) Correlation between CD8+ T cell infiltration and therapeutic response in YUMM1.7 tumors treated with BRAF/MEKi. (Simple linear regression) (J) Schematic illustrating the NK cytotoxicity assay. Created with BioRender.com. (K) Bar plot showing % NK cytotoxicity to YUMM1.7-GFP enriched from regressing tumors and residual disease at effector-to-target (E: T) ratios of 3:1 and 1:1. (n = 3 tumors, conducted in two independent experiments, multiple unpaired t-tests, mean ± SEM)

To determine if similar dynamics could be observed for CD8+ T cells, we examined the number of tumor-infiltrating CD8+ T cells during the treatment phases described above. The result showed that CD8+ T cells had a trend of increased abundance during tumor regression compared to growing tumors and remained at a similar level during residual disease (Fig. 4H and 4I). After tumors developed resistance and regrowth, the number of tumor-infiltrating CD8+ T cells decreased, but without statistical significance (Fig. 4H and 4I). We further analyzed whether CD8+T cells infiltrating in residual disease became more exhausted than in tumor regression. However, analysis of exhaustion markers, including Tim3, Tigit, Nkg2a, and PD-1, by flow cytometry did not show statistical differences (Fig.S6E-S6H). These findings align with previous observations that CD8+ T cell depletion does not significantly affect the onset of full resistance, likely due to unchanged infiltration and exhaustion status during these phases.

The scRNA-seq and flow cytometry analyses showed that tumor-infiltrating NK cells decreased in number but did not become more exhausted during therapy-tolerant residual disease compared to regressing tumors. However, it is unknown whether cancer cells in residual disease develop resistance to NK cell cytotoxicity. To determine this, we performed an NK cell cytotoxicity assay using primary NK cells co-cultured with melanoma cells isolated from regressing tumors and residual disease (Fig. 4J). YUMM1.7-GFP cells were FACS (fluorescence-activated cell sorting) isolated from regressing tumors and residual disease (Fig. S6I). Primary NK cells were enriched from naïve C57BL/6J mouse spleens using negative selection and stimulated with IL-2 and IL-15 (Fig. S6J). The NK cell cytotoxicity assay showed that residual disease had a similar susceptibility to NK cell cytotoxicity as regressing tumors at effector-to-target (E:T) ratios of 3:1 and 1:1 (Fig. 4K), indicating that therapy-tolerant residual disease did not develop resistance to NK cell cytotoxicity compared to regressing tumors. Together, these findings reveal that the transition from immune-inflamed to immune-excluded TME during BRAF/MEKi treatment is driven by the dynamics of NK cell infiltration rather than changes in CD8+ T cell abundance or exhaustion. Therapy-tolerant residual disease is characterized by significantly reduced NK cell infiltration but remains susceptible to NK cell cytotoxicity.

### Macrophages modulate NK cell infiltration during therapy-induced tumor regression

Since the tumor response to BRAF/MEKi is modulated by the amount of tumor-infiltrating NK cells and therapy-tolerant residual disease shows significantly less NK cell infiltration, we reasoned that increasing NK cell infiltration during residual disease could enhance the anti-tumor effect of BRAF/MEKi. Previous studies have shown that macrophages and NK cells can interact in tumors *(46–48)*. Our scRNA-seq analysis also indicated distinct macrophage populations between regressing tumors and residual disease (Fig. 2E). Aggregated cell-cell communication network analysis further revealed interactions between macrophages and NK cells in regressing tumors (Fig. S7A). Thus, we hypothesized that macrophages may create an immune microenvironment that favors NK cell infiltration during tumor regression.

To test our hypothesis, we first performed dual immunofluorescence staining of F4/80 and NK1.1 to examine the spatial interaction between macrophages and NK cells in the TME during both tumor regression and residual disease. Quantification of the number of macrophages near NK cells revealed a significant increase in macrophage-NK cell clusters in regressing tumors compared to residual disease (Fig. 5A and 5B), suggesting a potential interaction between macrophages and NK cells in regressing tumors.

**Figure 5.**
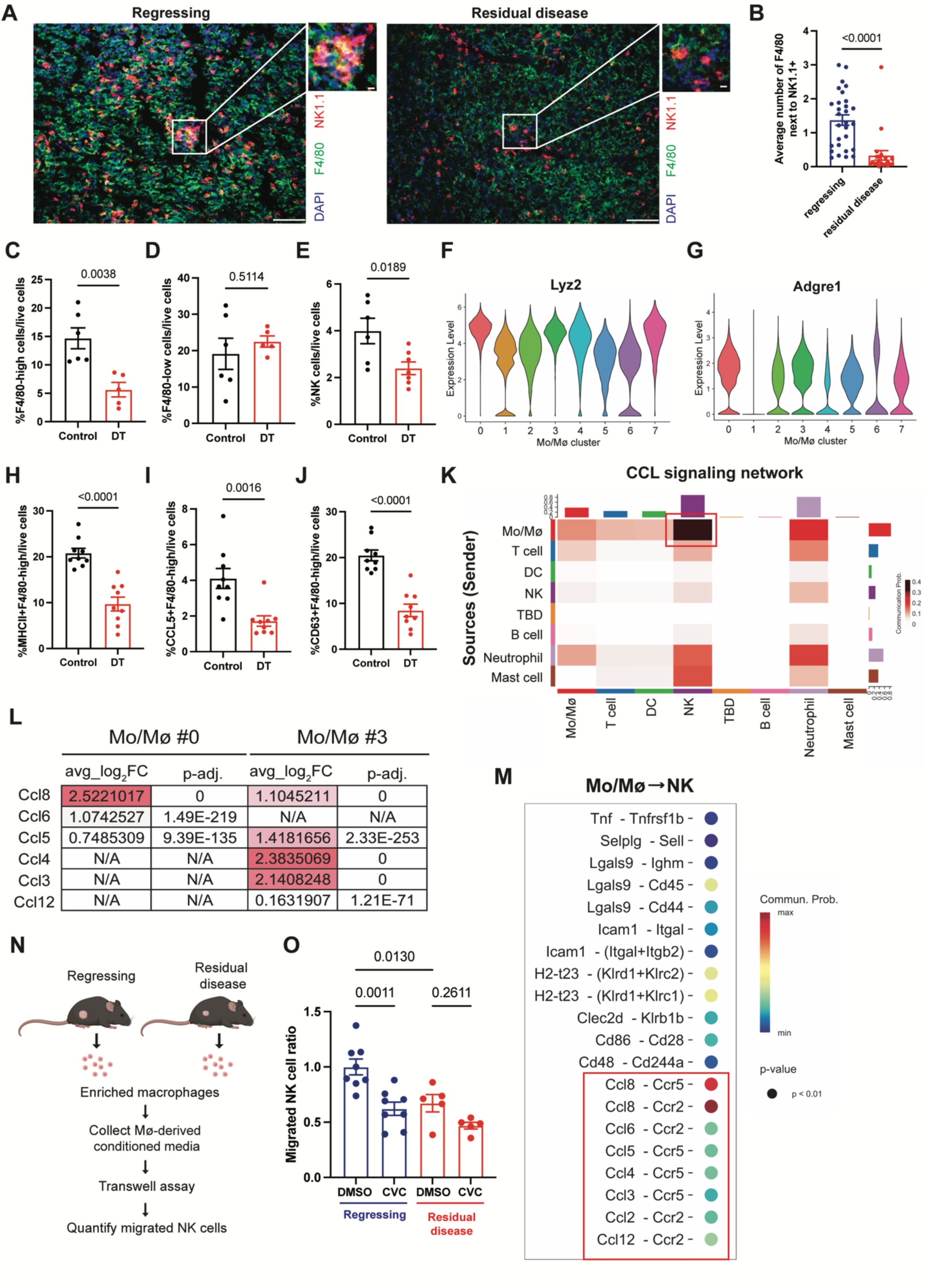
NK cell infiltration is modulated by specific macrophage populations via the CCR2/5 axis during tumor regression. (A and B) Representative immunofluorescence images of F4/80 (green), NK1.1 (red), and nuclear DAPI (blue) in regressing tumors and residual disease treated with BRAF/MEKi (A). Scale bar, 100 μm, 10 μm. Quantification of F4/80+ cells next to NK1.1+ cells (B). (n = 3 tumors with 4-11 total microscopy fields analyzed across all tissues, at least 1,600 counted NK1.1+ cells per tumor, two-tailed unpaired t-test, mean ± SEM) (C and D) Depletion efficiencies with diphtheria toxin (DT) (LysM-cre;iDTR) in tumors during tumor regression. Quantification of F4/80-high macrophages (C) and F4/80-low macrophages (D). (n = 5-6 tumors, two-tailed unpaired t-test, mean ± SEM) (E) Quantification of tumor-infiltrating NK cells in BRAF/MEKi-treated mice receiving diphtheria toxin (DT) (LysM-cre;iDTR) or control (NaCl in LysM-cre;iDTR or DT in WT mice) during tumor regression. (n = 6-7 tumors, two-tailed unpaired t-test, mean ± SEM) (F and G) Violin plot showing the expression of Lyz2 (F) and Adgre1 (G) in each macrophage cluster. Mo/Mϕ, monocyte/macrophage. (H—J) Quantification of MHC II+ (H), CCL5+ (I), or CD63+ (J) F4/80-high macrophages in BRAF/MEKi-treated mice receiving diphtheria toxin (DT) (LysM-cre;iDTR) or control (NaCl in LysM-cre;iDTR) during tumor regression. (n = 9 tumors, two-tailed unpaired t-test, mean ± SEM) (K) Heatmap depicting the CCL signaling network among different immune cell types during tumor regression, analyzed using CellChat. Mo/Mϕ, monocyte/macrophage. (L) Selected DEGs of multiple Ccl genes in macrophage Clusters 0 and 3. Mo/Mϕ, monocyte/macrophage. (M) Bubble plot showing ligand-receptor-based CellChat analysis during tumor regression, with macrophages as senders and NK cells as receivers. Mo/Mϕ, monocyte/macrophage. (N) Schematic illustrating experimental design for analysis of NK cells migrated toward macrophage-derived conditioned media. Mϕ, macrophage. Created with BioRender.com (O) Bar chart showing the migrated NK cell ratio in the transwell assay using conditioned media derived from macrophages isolated from regressing tumors or residual disease and primary splenocytes treated with a CCR2/5 inhibitor (CVC) or DMSO. CVC, cenicriviroc. (n = 5-8 tumors, conducted in two independent experiments, one-way ANOVA with Tukey’s multiple comparisons test, mean ± SEM)

To determine if macrophage presence affects relative NK cell recruitment, we injected clodronate liposomes with BRAF/MEKi for three days to deplete macrophages during the regressing phase (Fig. S7B). We validated the depletion efficiency by flow cytometry and confirmed that clodronate liposomes depleted circulating monocytes (both Ly6c-CD115+ and Ly6c+CD115+ monocytes) (Fig.S7C-S7E); however, clodronate liposomes only depleted <50% of the F4/80-high macrophages and did not reduce F4/80-low macrophages at all (Fig. S7F and S7G). The number of tumor-infiltrating NK cells also remained unchanged (Fig. S7H), possibly due to insufficient depletion of tissue-resident tumor-associated macrophages. To resolve this, we utilized the LysM-Cre; iDTR transgenic mouse model (Fig.S7I), which has been reported to exhibit rapid macrophage reduction in skin tissues after treatment with diphtheria toxin (DT) *(49)*. Using flow cytometry, we confirmed that a 3-day DT application led to a more profound depletion of F4/80-high macrophages than clodronate liposome injection, whereas F4/80-low macrophages remained unchanged (Fig. 5C and 5D). The reduction of F4/80-high macrophages resulted in a statistically significant decrease in tumor-infiltrating NK cells in regressing tumors (Fig. 5E), suggesting that F4/80-high macrophages in regressing tumors create a more favorable tumor immune microenvironment for NK cell infiltration.

To investigate if macrophages in residual disease had a similar impact on NK cell infiltration, we injected DT for three days in residual disease-bearing LysM-Cre;iDTR mice (Fig. S7J). Similarly, DT application significantly depleted F4/80-high macrophages, but not F4/80-low macrophages (Fig. S7K and S7L). However, NK cell infiltration was not affected by the depletion of F4/80-high macrophages in residual disease (Fig. S7M). Together, these results demonstrate the different functions of F4/80-high macrophages in tumor regression and residual disease, with F4/80-high macrophages in tumor regression playing a role in modulating NK cell infiltration. These results are consistent with our scRNA-seq analysis showing distinct macrophage phenotypes between tumor regression and residual disease (Fig. 2D-2F).

### Macrophage-NK cell crosstalk in therapy-induced tumor regression depends on CCR2/5 signaling

Given that increased NK cell infiltration is modulated by F4/80-high macrophages during tumor regression, we sought to identify the specific macrophage populations involved. To do this, we first analyzed our scRNA-seq data and found that Clusters 0 and 3 expressed the highest F4/80 (*Adgre1*) and LysM (*Lyz2*) expression among all macrophage clusters (Fig. 5F, 5G, S8A and S8B), suggesting that DT application may selectively deplete these two populations. Clusters 0 and 3 were the most abundant macrophage populations and exclusively existed in regressing tumors (Fig. 2D, 2E, S2B and S2C). Using spectral flow cytometry, we validated a variety of markers that were upregulated in Clusters 0 and 3 in our scRNA-seq data, including F4/80, Ccl5, Cd63, and MHC II (Fig. S8C). The result showed that DT selectively depleted macrophages expressing these markers in regressing tumors (Fig. 5C, 5H-5J, and S8D), indicating that these specific macrophage populations, which are present during tumor regression, modulate NK cell infiltration.

Next, we set out to determine how macrophages and NK cells interact with each other in regressing tumors. Cellchat analysis of our scRNA-seq data showed a CCL signaling network sending from macrophages to NK cells with the highest probability in regressing tumors (Fig. 5K). In addition, Clusters 0 and 3 also had significantly higher expression of Ccl ligands (Fig. 5L and S8E). Ligand-receptor analysis further indicated that macrophages could use multiple CCL ligands, including Ccl2, Ccl3, Ccl4, Ccl5, Ccl6, Ccl8, and Ccl12, to interact with Ccr2/Ccr5, which are expressed on NK cells in regressing tumors (Fig. 5M). To validate whether the recruitment of NK cells by macrophages is dependent on the CCR2/5 axis during tumor regression, we isolated macrophages from regressing tumors and residual disease and collected macrophage-derived conditioned media from both groups. We then performed a transwell assay using macrophage-derived conditioned media and primary splenocytes treated with cenicriviroc (CVC, a CCR2/5 inhibitor) or vehicle control, and then quantified migrated NK cells by flow cytometry (Fig. 5N and S8F). The result showed that conditioned media derived from macrophages isolated from regressing tumors significantly enhanced NK cell migration compared to conditioned media from macrophages isolated from residual disease (Fig. 5O). With CCR2/5 inhibition, NK cell migration toward macrophage-derived conditioned media was abolished, and this effect was more pronounced in the regressing tumor group (Fig. 5O, S8G and S8H). Collectively, these results demonstrate that specific macrophage populations in tumor regression regulate NK cell infiltration via the CCR2/5 axis.

### Reprogramming macrophages and enhancing NK cell infiltration shifts the TME from immune-excluded to -inflamed, improving targeted therapy efficacy

Given that NK cell infiltration is regulated by specific macrophage populations, we sought to identify a strategy to reprogram macrophages and enhance NK cell infiltration, with the goal of converting the immune-excluded TME into an immune-inflamed state to better support anti-tumor immunity. Previous studies have shown that protein tyrosine phosphatase non-receptor type 22 (Ptpn22) is a key regulator of T cell receptor (TCR) signaling, and a lack of Ptpn22 may enhance the antitumor immune response *(50–52)*. Additionally, inhibition of Ptpn22 has been reported to polarize macrophages to an M1-like state *(53, 54)*. Although we are not aware of any studies describing a function for Ptpn22 in NK cells, a recent report demonstrated that Ptpn22 inhibition can increase NK cell infiltration in a colorectal cancer mouse model *(54)*.

Our scRNA-seq data revealed a high expression level of Ptpn22 in T cells and NK cells, and a moderate expression level in macrophages, suggesting that Ptpn22 might be a potential target in our model (Fig. 6A). To test this hypothesis, we examined whether Ptpn22 inhibition could enhance NK cell migration. Splenic NK cells treated with L-1 *(54)*, a Ptpn22 inhibitor, showed significantly increased migration in a transwell assay compared to DMSO-treated controls (Fig. S9A, S9B). Next, we treated YUMM1.7-bearing mice with L-1 starting on day 10 of BRAF/MEKi treatment, during the establishment of residual disease (Fig. 6B). L-1 treatment significantly increased the number of tumor-infiltrating NK cells (Fig. 6C and 6D), with a trend of increased granzyme B expression, indicating enhanced activation of NK cells (Fig. S9C). Moreover, L-1 treatment led to a marked increase in F4/80-high macrophages, while the population of F4/80-low macrophages remained unchanged (Fig. 6E and 6F). Ccl5+ and Cd63+ F4/80-high macrophages also increased, and there was a trend of increased MHC II+ F4/80-high macrophages in L-1 treated tumors (Fig. 6G-6I). These findings demonstrate that L-1 treatment alters the residual tumor immune microenvironment, shifting it toward a more regression-like, immune-inflamed phenotype. This was marked by a heightened infiltration of NK cells and a predominance of regression-like macrophages, particularly an increase in F4/80-high macrophages, with amplified expression of Ccl5, Cd63 and MHC II (Fig. 4B, 4D, 6J-6N).

**Figure 6.**
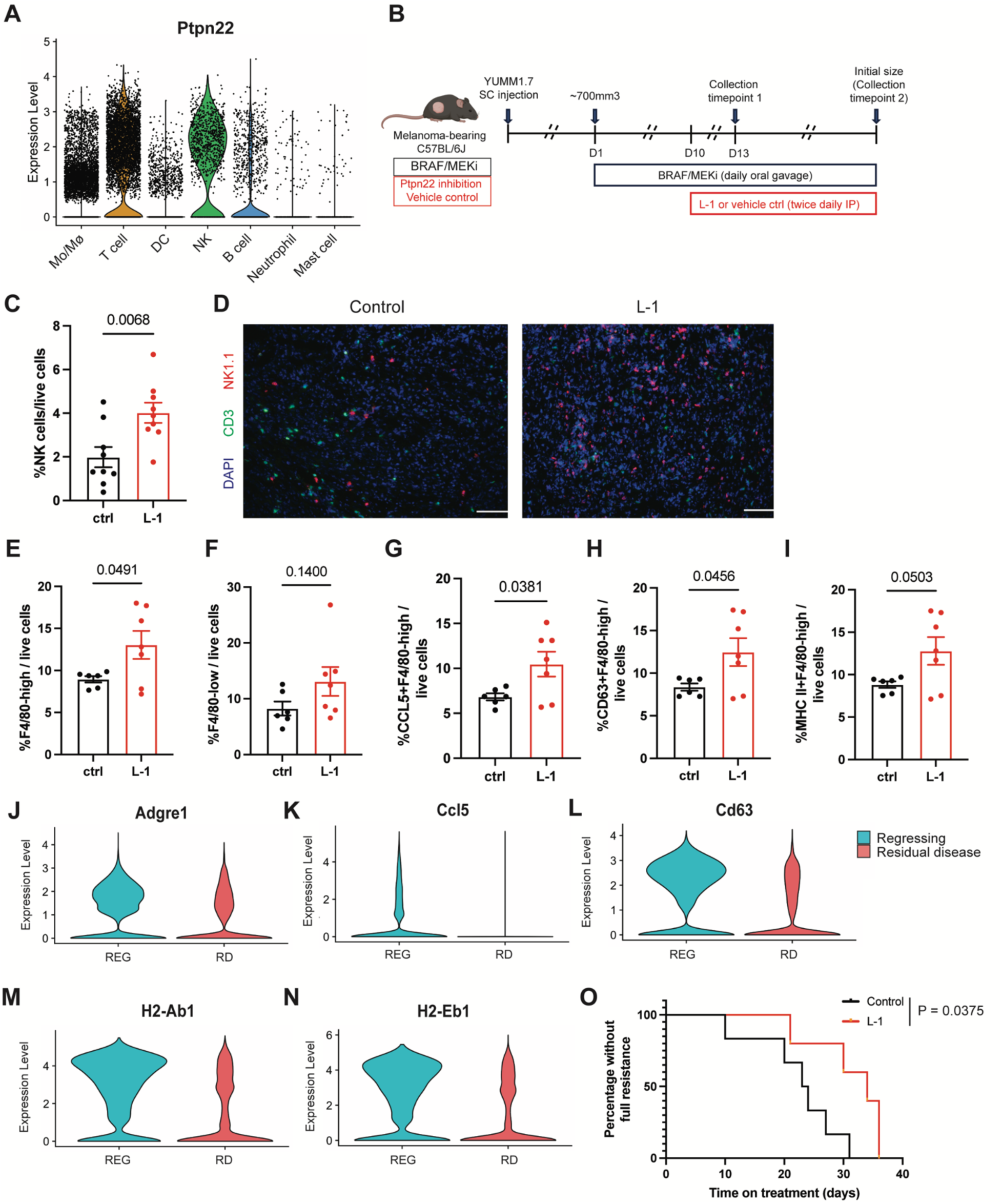
Ptpn22 inhibition increases infiltration of NK cells and regression-like macrophages, resulting in enhanced tumor control by BRAF/MEKi. (A) Violin plot showing Ptpn22 expression in each immune cell type. Mo/Mϕ, monocyte/macrophage. (B) Schematic illustrating experimental design for targeting Ptpn22 in YUMM1.7-bearing mice. Created with BioRender.com. (C) Quantification of tumor-infiltrating NK cells in BRAF/MEKi-treated mice receiving L-1 or vehicle control on day 13. (n = 9 tumors, two-tailed unpaired t-test, mean ± SEM) (D) Representative immunofluorescence images of NK1.1 (red), CD3 (green) and nuclear DAPI (blue) of BRAF/MEKi-treated tumors in the indicated treatment groups on day 13. NK cells are NK1.1+CD3-. Scale bar 100 μm. (E and F) Quantification of tumor-infiltrating F4/80-high (E) or F4/80-low (F) macrophages in BRAF/MEKi-treated mice receiving L-1 or vehicle control on day 13. (n = 6-7 tumors, two-tailed unpaired t-test, mean ± SEM) (G—I) Quantification of tumor-infiltrating Ccl5+ (G), Cd63+ (H), or MHC II+ (I) F4/80-high macrophages in BRAF/MEKi-treated mice receiving L-1 or vehicle control on day 13. (n = 6-7 tumors, two-tailed unpaired t-test, mean ± SEM) (J—N) Violin plot showing the expression of Adgre1 (J), Ccl5 (K), Cd63 (L), and H2-Ab1/Eb1 (M and N) in the monocyte/macrophage population between regressing tumors and residual disease. REG, regressing; RD, residual disease. (O) Kaplan-Meier curve for BRAF/MEKi-treated mice bearing YUMM1.7 receiving L-1 or vehicle control. (n = 5-6 tumors, Log-rank Mantel-Cox test)

Since a previous study reported that L-1 treatment increased the infiltration and activation of CD8+ T cells in the MC38 colorectal tumor model *(54)*, we also quantified the tumor-infiltrating CD8+ T cells and granzyme B expression levels; however, we did not observe a significant difference (Fig. S9D and S9E).

After confirming that L-1 treatment reprograms macrophage populations and increases NK cell infiltration in our model, we further tested our hypothesis that converting an immune-excluded TME to an immune-inflamed state by manipulating macrophages and NK cells in drug-tolerant residual disease can enhance the anti-tumor efficacy of targeted therapy. To do this, we added L-1 from day ten after starting BRAF/MEKi treatment and evaluated the time required for YUMM1.7-bearing mice to relapse (Fig. 6B). As expected, Ptpn22 inhibition significantly delayed the onset of full resistance to BRAF/MEKi by altering macrophages and improving NK cell infiltration during residual disease (Fig. 6O, S9F, and S9G). To determine if the enhanced anti-tumor effect caused by L-1 is dependent on NK cells, we evaluated the time required for YUMM1.7-bearing mice treated with L-1 to achieve full resistance to BRAF/MEKi after NK cell depletion (Fig. S9H). Depletion of NK cells completely abrogated the effect of L-1, indicating the necessary role of NK cells in L-1-treated tumors (Fig. S9I).

In summary, these findings demonstrate that reprogramming macrophage phenotypes and enhancing NK cell infiltration can transform an immune-excluded tumor microenvironment into an immune-inflamed state, significantly amplifying the anti-tumor effects of BRAF/MEKi. This reveals a powerful strategy to overcome resistance by harnessing innate immunity to reinforce and sustain therapeutic efficacy.

### Infiltration of NK cells and specific macrophage populations positively correlate with the response to BRAF/MEKi in patients with melanoma and tyrosine kinase inhibitors in lung cancer

To determine whether a similar tumor immune microenvironment exists in human melanomas treated with BRAF/MEKi, we analyzed a publicly available scRNA-seq dataset that sequenced fine-needle aspirates (FNA) from four melanoma patients before and at different time points following BRAF/MEKi treatment *(55)*. Patients 1 and 2 were both resistant to BRAF/MEKi. Patient 3 consistently responded to BRAF/MEKi at all biopsy time points, whereas Patient 4 initially responded to BRAF/MEKi but later developed resistance (Fig. 7A). The UMAP plots displaying different cell types in each patient are shown in Figure 7B.

**Figure 7.**
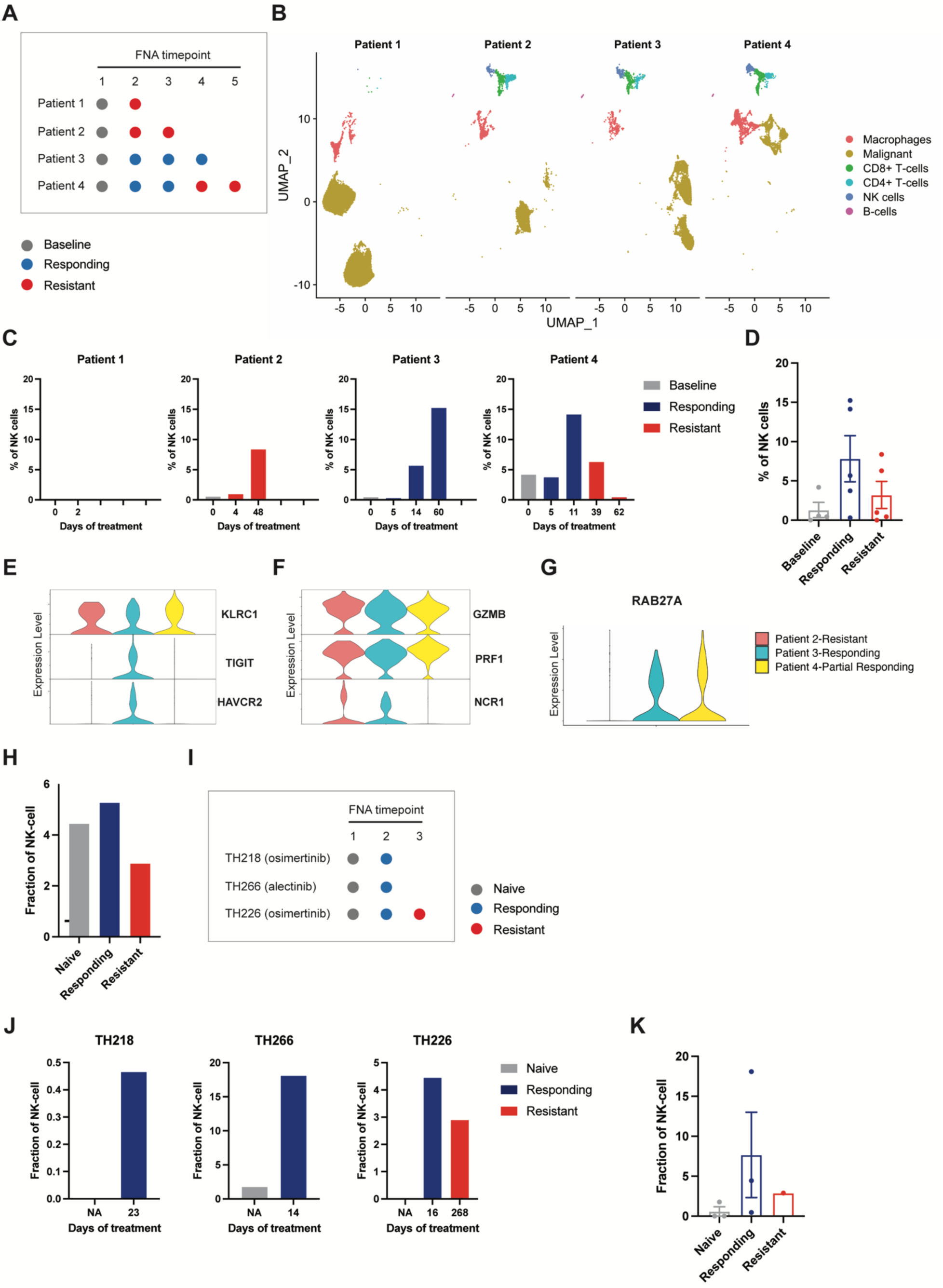
NK cell infiltration positively correlates with the targeted therapy response in human cancers. (A) Schematic illustrating the timeline and response to BRAF/MEKi for four melanoma patients in GSE229908. Each dot represents one FNA biopsy. Grey represents baseline (before BRAF/MEKi), blue represents responding tumors, and red represents BRAF/MEKi-resistant tumors. (B) UMAP plot showing the cell populations of the four melanoma patients. (C) Bar plots showing NK cell quantification at each FNA timepoint in four melanoma patients. (D) Quantification of NK cells in FNA biopsies grouped into Baseline (before BRAF/MEKi treatment), Responding, and Resistant groups. (mean ± SEM) (E—G) Violin plots showing the expression of exhaustion markers (E), activation markers (F), or RAB27A (G) in the NK cell population in Patient 2 (Resistant), 3 (Responding) and 4 (Partial Responding). (H) Bar plot showing NK cell fractional changes across three treatment states in PRJNA591860, containing 49 lung cancer biopsies before and during TKI treatment. (I) Schematic illustrating the timeline and response to TKI treatment in three lung cancer patients with sequential biopsies in PRJNA591860. Each dot represents one FNA biopsy. Grey represents naïve (before TKI treatment), blue represents responding tumors, and red represents resistant tumors. (J) Bar plots showing NK cell fractional changes at each FNA timepoint in three lung cancer patients. (K) NK fractional changes in FNA biopsies from the three patients grouped into Naïve, Responding, and Resistant groups. (mean ± SEM)

To assess if NK cell infiltration is positively correlated with tumor response to BRAF/MEKi in patient samples, we quantified the percentage of NK cells among all sequenced cells (Fig. 7C). Notably, Patient 1, who was resistant to BRAF/MEKi, had almost no NK cells. Patient 4, consisting of responding and resistant biopsies, was most similar to the BRAF/MEKi tumor response in our mouse model. The percentage of NK cells in this patient demonstrated a positive correlation with the tumor response to BRAF/MEKi (i.e., NK cell infiltration increased in responding biopsies and decreased in resistant biopsies), which is consistent with our findings (Fig. 4A, 4B and 7C). Next, we grouped all the biopsy timepoints based on the tumor response, including baseline (i.e., before treatment), responding, and resistant from each patient. We observed that the number of tumor-infiltrating NK cells increased in responding tumors and decreased in resistant tumors (Fig. 7D). The average percentage of tumor-infiltrating NK cells in this human dataset was similar to that observed in our *in vivo* preclinical analysis, thus validating our findings (Fig. 4B, 4C and 7D).

To assess if tumor-infiltrating NK cells exhibit different phenotypes between responding and resistant tumors, we analyzed multiple inhibitory receptors (i.e., *KLRC1*, *TIGIT*, and *HAVCR2*) and activating molecules (i.e., *GZMB*, *PRF1*, and *NCR1*) in sequenced NK cells from Patients 2, 3, and 4 (Patient 1 had no detected NK cells, and thus was excluded). The results indicated that tumor-infiltrating NK cells did not exhibit increased exhaustion in resistant tumors compared to responding and partially responding tumors (Fig. 7E). In addition, the expression of activating receptors did not decrease in NK cells infiltrating resistant tumors (Fig. 7F). These results are consistent with our analysis of the mouse model (Fig. 4E-4G). Notably, NK cells from Patient 2 exhibited decreased expression of RAB27A, a critical gene that controls NK cell cytotoxicity (Fig. 7G). This observation suggests that NK cells from Patient 2 have a dysfunctional phenotype in resistant tumors and resemble NK Cluster 0, which was the predominant NK cluster in therapy-tolerant residual disease in our mouse model (Fig. 2L-2O). The analysis of NK cells in these patient melanoma scRNA-seq datasets supports our findings. This suggests that our mouse model replicates the NK cell infiltration observed in melanoma patients treated with BRAF/MEKi, and indicates that NK cells significantly impact the tumor response to BRAF/MEKi in both contexts.

In addition to the NK cell profile, we also interrogated the macrophage populations in these patient samples. To determine whether the presence of Clusters 0 and 3 identified in our mouse model correlates with the response to BRAF/MEKi in patient melanomas, we compared the macrophage signature of Clusters 0 and 3 to melanoma patient macrophage populations. Although patient-specific heterogeneity was observed according to the macrophage UMAP (Fig. S10A), the results revealed that Patient 3 (responding) had a signature that most closely resembled Clusters 0 and 3 (Fig. S10B and S10C).

In summary, we identified that NK cell infiltration is associated with tumor response to BRAF/MEKi in melanoma patients using publicly available data. Similar to the NK cells infiltrating mouse tumors, the NK cells in patient melanomas did not become more exhausted in resistant biopsies compared to responsive biopsies, emphasizing the importance of increasing NK cell infiltration rather than relying only on checkpoint blockade to improve the anti-tumor effect of BRAF/MEKi.

To investigate whether our findings could extend to targeted therapies beyond BRAF/MEKi, we analyzed a publicly available scRNA-seq dataset consisting of 49 clinical biopsies from 30 advanced-stage lung cancer patients, obtained before and during tyrosinase kinase inhibitor (TKI) treatment, including EGFR, ALK, BRAF, and MEK inhibitors *(56)*. The NK cell population was overlooked in the original analysis in the published paper. By re-analyzing the dataset, we identified an NK cell population using human NK cell markers, including KLRD1, KLRF1, NCAM1, and FCGR3A, while excluding CD3 gene expression (Fig. S11A and S11B).

We compared the immune cell composition across three treatment phases: naïve (before treatment), responding (biopsies obtained during tumor regression or stable disease as determined by clinical imaging), and resistant (subsequent progressive disease following acquired drug resistance). Consistent with our preclinical models and findings in human melanoma patients, NK cell fractional change was positively correlated with treatment response, showing an increase during the response phase compared to the naïve and resistant phases (Fig. 7H).

Obtaining consecutive clinical tumor biopsies from advanced-stage lung cancer patients during treatment is inherently challenging, as most tumors regress by 50% or more during targeted therapy *(56)*. Despite this limitation, three patients (TH218, TH266, and TH226) in the dataset had matched tumor biopsies collected at multiple time points, with patient TH226 providing samples across all three phases (naïve, responding, and resistant) (Fig. 7I). Analysis of these cases revealed a consistent pattern: NK cell fractions were higher in the responding biopsies than in the naïve and resistant biopsies (Fig. 7J and 7K). These findings suggest that NK cells serve as determinants of treatment efficacy in therapies targeting diverse aberrant signaling pathways, including TKIs, in lung cancer. This highlights the broader applicability of our findings across different tumor types and targeted therapies.

## DISCUSSION

While immune-inflamed and immune-excluded tumors exhibit distinct responses to cancer therapies *(24, 57)*, it is unclear whether the pre-existing immune TME persists throughout treatment or undergoes dynamic changes. Additionally, while adaptive immunity has been the primary focus in cancer therapy, emerging evidence has highlighted the critical role of innate immunity *(25–27)*. Here, we reveal a previously unrecognized transition from a targeted therapy-induced immune-inflamed TME to an immune-excluded state as tumors acquire resistance, marked by reduced NK cell infiltration and altered macrophage phenotypes. We demonstrate that BRAF/MEKi therapeutic efficacy in melanoma critically depends on tumor-infiltrating NK cells, and that therapy-tolerant residual disease remains susceptible to NK cell cytotoxicity. Moreover, we identify macrophage-NK cell crosstalk via the CCR2/5 axis as a key regulator of NK cell infiltration during residual disease. Modulating this interaction by inhibiting Ptpn22 restores the immune-inflamed phenotype, enhances NK cell infiltration, and improves BRAF/MEKi efficacy. Finally, we validate these findings in melanoma and lung cancer patients, confirming that NK cell infiltration correlates with response to targeted therapies.

We observe robust NK cell infiltration during therapy-induced tumor regression, followed by a decline in therapy-tolerant residual disease. While NK cells are often considered ineffective in solid tumors due to poor infiltration and immunosuppressive factors *(46)*, our findings demonstrate their ability to infiltrate tumors during regression and serve as critical determinants of therapy efficacy. Additionally, we identify a specific tumor-associated macrophage (TAM) population marked by elevated F4/80, Ccl5, MHC II, and Cd63 expression, which contributes to an anticancer milieu that promotes NK cell infiltration and enhances BRAF/MEKi effectiveness. Unlike previously reported TAMs that support NK cell cytotoxicity via IL-12, IL-15, IL-18, and TNF-α *(46, 48, 58)*, this subset appears to drive NK cell recruitment primarily through the CCR2/5 signaling pathway. Future studies should investigate whether these macrophages directly contribute to tumor cell death and identify the factors that polarize them toward an antitumor phenotype. Given the plasticity and high abundance of TAMs, targeting macrophage polarization represents a promising strategy to enhance cancer therapy efficacy by reprogramming the TME.

We further demonstrate that innate immune components, specifically NK cells and TAMs, can be harnessed to convert an immune-excluded TME into an immune-inflamed state—using Ptpn22 inhibition as a proof-of-concept approach—resulting in improved BRAF/MEKi tumor control. To the best of our knowledge, no prior studies have examined the synergistic effects of Ptpn22 inhibition and targeted therapy. Notably, we provide the first evidence that the therapeutic effect of Ptpn22 inhibition is dependent on NK cells rather than CD8⁺ T cells, as previously reported *(54)*. Although beyond the scope of this study, the precise mechanisms by which Ptpn22 inhibition influences macrophage reprogramming and NK cell function remain unclear and warrant further investigation. Additionally, future studies should explore factors beyond TAMs that regulate NK cell infiltration to develop broader strategies for transforming immune-excluded TMEs into inflamed states by harnessing innate immune components. Ultimately, unlocking innate immune-mediated anti-tumor responses will synergize with adaptive immunity, generating a potent and coordinated anti-tumor response to overcome therapy resistance.

Finally, we validated our preclinical findings using two independent patient scRNA-seq datasets, including melanoma patients treated with BRAF/MEKi and lung cancer patients treated with TKIs. Consistent with our preclinical models, NK cell infiltration positively correlated with treatment efficacy, reinforcing the broader clinical relevance of NK cells in targeted therapy responses. However, the correlation between macrophages and treatment response was less pronounced. This discrepancy may stem from differences between mouse and human macrophages, as highlighted by previous and recent studies *(55, 59)*, or the limited sample size of the patient datasets analyzed. A constraint of our study is the challenge of assessing NK cell-macrophage interactions in available human datasets. Unlike preclinical models, where these interactions can be rigorously studied, the lack of large, well-annotated human cancer datasets with sufficient sample sizes and time-point-specific immune profiling limits such analyses. Future studies will require larger patient cohorts and comprehensive immune profiling to better elucidate the interactions between NK cells and macrophages and their contributions to therapeutic responses. Together, these findings suggest that the relationship between NK cell infiltration and therapeutic response in BRAF/MEKi-treated melanoma may extend to other cancers and targeted therapies, including TKIs for lung cancer. This underscores the broader relevance of enhancing NK cell infiltration as a strategy to improve targeted therapy efficacy across different tumor types.

In conclusion, we demonstrate that therapy-tolerant residual disease exhibits an immune-excluded TME that is not pre-existing but emerges from an immune-inflamed state during tumor regression. This transition is driven by loss of NK cell infiltration and macrophage reprogramming. Although targeted therapies are designed to inhibit aberrant kinase signaling in tumor cells, unlike immunotherapies, which directly modulate immune cells, our findings reveal that innate immunity plays a critical role in their efficacy. Crucially, we show that modulating innate immune components can reprogram the TME to restore an immune-inflamed state, enhancing targeted therapy efficacy. These findings emphasize the essential role of innate immunity in optimizing targeted therapy responses, and highlight a new therapeutic avenue for overcoming resistance.

## MATERIALS AND METHODS

### Study design

This study aimed to comprehensively characterize the dynamic evolution of the tumor immune microenvironment throughout different phases of targeted therapy and to identify key determinants that can be modulated to shift an immune-excluded TME toward an immune-inflamed state, thereby improving therapeutic efficacy. To deconvolute the immunological landscape during tumor regression and drug-tolerant residual disease, we performed scRNA-seq of immune cells isolated from BRAF/MEKi-induced regressing tumors and drug-tolerant residual disease (three biological replicates per group). Immune cell depletion experiments were conducted in BRAF/MEKi-treated tumor-bearing mice to determine which immune cell populations are essential for an optimal tumor response. Flow cytometry and IF staining were used to validate scRNA-seq findings and confirm depletion efficiency. To investigate NK cell-macrophage crosstalk, we employed scRNA-seq analysis, macrophage depletion, flow cytometry, IF staining, and transwell assays. To assess whether modulating innate immunity could enhance targeted therapy response, Ptpn22 inhibition was performed in BRAF/MEKi-treated melanoma-bearing mice, followed by immune cell infiltration/reprogramming analysis and depletion studies to determine the role of specific immune cells in tumor control. Human patient data from two independent publicly available datasets, including melanoma patients treated with BRAF/MEKi and lung cancer patients treated with TKI, were analyzed for clinical relevance. For preclinical experiments, mice were randomized into different treatment groups. The endpoint of the mice for assessing time to resistance was when tumors reached the initial size before BRAF/MEKi treatment (i.e. ∼700mm^3^ in size). The endpoint for regressing tumors and growing tumors was after 3 days of BRAF/MEKi or vehicle control treatment. The endpoint for residual disease was after 14 days of BRAF/MEKi treatment.

### Tumor cell lines

YUMM1.7 (ATCC), YUMM1.7-GFP (gift from Dr. Amanda Lund, New York University), and YUMMUV1.7 and YUMMUV3.3 (gift from Dr. Brian Gabrielli, The University of Queensland) were cultured in DMEM/F-12 containing 10% FBS, 1% non-essential amino acids and penicillin/streptomycin in a 5% CO2 cell culture incubator at 37°C. YUMM1.7 cells tested negative for mycoplasma using a PCR-based method.

### Mouse models

C57BL/6J and NSG mice were purchased from the Jackson Laboratory and bred in-house. *LysM-Cre* (Jax, stock #004781) and *Rosa26-lsl-iDtr* (Jax, stock #007900) were purchased from the Jackson Laboratory and bred in-house. For syngeneic immunocompetent melanoma mouse models, age-matched 7- to 12-week-old mice were injected with YUMM1.7, YUMM1.7-GFP, YUMMUV1.7, or YUMMUV3.3 (1 × 10^6^) subcutaneously into the right and left flanks. Tumor volume was measured daily using a digital caliper (V =W^2^ × L/2). All mouse treatments were approved by the Institutional Animal Care and Use Committee (IACUC) at Cornell University under protocol number 2016-0034. Mice were maintained under pathogen-free conditions at the Cornell University College of Veterinary Medicine by the Center of Animal Resources and Education (CARE). To assess resistance onset in YUMM1.7-bearing NSG mice, in YUMM1.7 and YUMMUV1.7-bearing C57BL/6J mice during NK cell depletion and in *LysM-cre; iDTR* mice during macrophage depletion in regressing tumors, both male and female mice were used. Only male mice were used in all other experiments.

### Targeted therapy treatment

Mice received BRAF/MEKi therapy at a dose of 25 mg/kg dabrafenib (Medchem Express, catalog no. HY-14660) and 0.15 mg/kg trametinib (Medchem Express, catalog no. HY-10999). Dabrafenib and trametinib were initially dissolved in dimethyl sulfoxide (DMSO) and subsequently diluted in carboxymethylcellulose (CMC; 0.5% w/v) and Tween 80 (0.05% v/v) in sterile PBS to a final volume of 200 μl, and administered to mice daily by oral gavage. Mice bearing melanoma tumors were treated with BRAF/MEKi when tumors reached ∼700mm^3^.

### CD45+ cell enrichment

Tumors were collected and dissociated in DMEM/F12 media containing collagenase Type I (20 mg/ml) (Worthington, catalog no. LS004196) at 37°C for 75 minutes. Digested materials were filtered through 40 μm cell strainers to obtain a single-cell suspension and then washed with MACS buffer (PBS supplemented with 0.5% BSA + 1mM EDTA). Cells were counted using a hemocytometer. Single-cell suspensions were incubated with CD45 microbeads (Miltenyi Biotec, catalog no. 130-052-301) and then passed through MS Columns (Miltenyi Biotec, catalog no. 130-042-201), according to the manufacturer’s instructions. Enriched cells were collected and underwent downstream experiments.

### Single-cell RNA sequencing and analysis

Single-cell suspensions were obtained, followed by CD45+ cell enrichment as described above and subsequently submitted to the Cornell University BRC Genomics Core Facility (RRID: SCR_021727), where they were run on a Chromium X instrument and libraries were prepared following the Chromium Next GEM Single Cell 3’ High Throughput RNA-Seq Assay v3.1 Dual Index user guide (10x Genomics, CG000416, RevC). Sequencing results were processed with 10x Genomics Cell Ranger using the mouse (GRCm39) reference. All downstream analyses were performed using Seurat (v5.1.0). Raw data were merged and QC was analyzed with nFeature_RNA > 200 and nFeature_RNA < 8000. Contaminated non-CD45+ cells (e.g., fibroblasts and cancer cells) were excluded, resulting in a final dataset of 41,588 immune cells. For data integration and scaling, the functions “IntegrateData” and “ScaleData” were used. For the visualization of cells in a two-dimensional space, highly variable genes were selected for principal component analysis (PCA), and the first 50 PCs were used for clustering analysis and visualization of the clustering results by Uniform Manifold Approximation and Projection (UMAP). Differential gene expression analysis was performed using the “FindAllMarkers” function (avg_log2FC > 1). The top differentially expressed genes per cluster based on log2FC were plotted in a heatmap using the “DoHeatmap” function, and selected genes of interest were visualized in a dot plot using the “DotPlot” function. Macrophage biological process scoring was calculated using the “AddModuleScore” function from the signature reported in the previous studies *(36, 37)*. Cell-cell communication and ligand-receptor interactions were analyzed using the “CellChat” package.

The publicly available scRNA-seq dataset of melanoma patient FNA biopsies from Vasilevska et al. *(55)* was downloaded from GEO (GSE229908). The downloaded RDS file is converted into a Seurat object using R. All downstream analyses were performed using Seurat (v5.1.0). TTS/TTR macrophages in the metadata were combined as one macrophage population, and all cell types were visualized between different patients on UMAP. The signature of macrophage Clusters 0 and 3 identified in our mouse scRNA-seq dataset was applied to the human data using the “AddModuleScore” function. Violin plots were generated using the “VlnPlot” function to compare the enrichment scores of these clusters.

The publicly available scRNA-seq dataset of patient lung cancer biopsies from Maynard et al.*(56)* was downloaded from Google Drive on GitHub at Czbiohub/scell_lung_adenocarcinoma. The downloaded RData file of immune cells was analyzed using Seurat (v5.1.0) in R. The T cell cluster in the metadata was re-annotated to identify the NK cell population.

### Macrophage depletion by clodronate liposomes

For short-term (< one week) macrophage depletion in regressing tumors, melanoma-bearing C57BL/6J mice were injected intravenously daily with 150 μl (per 20g mouse weight) of clodronate liposomes (CL) or PBS liposomes (PL) (Encapsula NanoSciences, catalog no. CLD-8901) as a control for three days from the first day of BRAF/MEKi treatment (days 1 to 3). For long-term macrophage depletion to assess resistance onset, melanoma-bearing C57BL/6J mice were injected intravenously every other day with 150 μl (per 20g mouse weight) CL or PL from day 10 for five injections (days 10 to 18) and then 100 μl (per 20g mouse weight) of CL or PL twice a week until endpoint; lower dosage and frequency were administered due to increased risk of infection resulting from long-term depletion.

### Macrophage depletion by diphtheria toxin

Diphtheria toxin (DT) (Sigma, catalog no. D0564) was dissolved in ddH2O at a stock concentration of 600 ng/μL and stored at −80 °C. Before use, the stock solution was diluted with sterile 0.9% NaCl saline to a final concentration of 2.52 ng/μl with a total volume of 200 μl, which was administered via intraperitoneal injection. To deplete macrophages during tumor regression, melanoma-bearing LysM-cre; iDTR mice received daily injections from day 1 to day 3 of BRAF/MEKi treatment. To deplete macrophages during the residual disease phase, injections were administered daily from day 10 to day 12 of treatment. The control group included *LysM-cre; iDTR* mice injected with an equivalent amount of 0.9% NaCl saline and WT mice injected with DT.

### CD8+ T cell depletion

C57BL/6J mice were intraperitoneally injected with an anti-CD8b antibody (200 μg) (clone Ly-3.2, BioXcell, catalog no. BE0223) three days and one day prior to tumor transplantation and with continued injection once a week until tumor collection. Control mice were injected with an equivalent amount of IgG1 isotype control (BioXcell, catalog no. BE0088).

### NK cell depletion

C57BL/6J mice received intraperitoneal injections of anti-asialo GM1 (ASGM1) antibody (Wako Pure Chemicals, catalog no. 986-10001) (20 μl) in sterile PBS two days prior to tumor transplantation or L-1 treatment, with subsequent injections every four days until tumor collection. Control mice were administered rabbit serum (20 μl) in sterile PBS.

### Flow cytometry and cell sorting

Tumors were collected and dissociated in DMEM/F12 media containing collagenase Type I (20 mg/ml) (Worthington, catalog no. LS004196) at 37°C for 75 minutes. Digested materials were filtered through 70 μm cell strainers to obtain a single-cell suspension. Cells were counted using a Scepter 3.0 Handheld Automated Cell Counter (Millipore Sigma, catalog no. PHCC340KIT). Cell suspensions were incubated with an anti-CD16/32 Fc receptor blocker (BioLegend, catalog no. 156603) to block mouse Fc receptors, and the Zombie UV Fixable Viability Kit (BioLegend, catalog no. 423107) or the LIVE/DEAD Fixable Violet Dead Cell Stain Kit (Thermo Fisher, catalog no. L34964) to exclude dead cells in PBS for 15 minutes at room temperature before surface staining. Surface staining was performed in MACS buffer or Brillant Staining Buffer (BD, catalog no. 563794) for 30 minutes at 4°C (antibodies are listed in Table S1). For intracellular staining, cells stained with surface markers were fixed and permeabilized using the Intracellular Fixation & Permeabilization Buffer Set (Thermo Fisher, catalog no. 88-8824-00), according to the manufacturer’s instructions, followed by staining with antibodies (Table S1). Following intracellular staining, cells were resuspended in MACS buffer, and data were acquired on a BD FACSymphony A3 or BD FACSymphony A5. Compensation, analysis, and visualization of the flow cytometry data were performed using FlowJo Software (RRID: SCR_008520). For the NK cell cytotoxicity assay, YUMM1.7-GFP cells were sorted from regressing tumors and residual disease using a BD FACSMelody. Dead cells were excluded using the LIVE/DEAD Fixable Violet Dead Cell Stain Kit.

### Immunofluorescence (IF) staining

Tumors were collected and embedded in optimal cutting temperature (OCT) compound (Thermo Fisher Scientific, catalog no. 23730571), snap-frozen, and stored at −80°C. For IF staining of frozen tissue, 8-μm sections of tumors were cut on a cryostat, fixed in 10% neutral-buffered formalin for 10 minutes, and washed twice with distilled water for 10 minutes each. Subsequently, each section was blocked for non-specific protein interactions with 10% donkey serum in PBST, followed by overnight staining with primary antibodies at 4°C (Table S1). The following day, slides were washed with PBST and incubated for 1 hour at room temperature with secondary antibodies conjugated to fluorochromes. Fluoroshield mounting medium with DAPI (Abcam, catalog no. ab104139) was used for nuclei staining. Images were captured using a Leica DM2500 upright microscope equipped with a DFC7000T camera. Quantification of macrophages adjacent to NK cells was performed using CellProfiler (RRID: SCR_007358). F4/80+ cells within a neighbor distance of 5 pixels (equivalent to 4.46 μm) from NK1.1+ cells were identified and counted. The quantification analyses were performed by a person who was unaware of the experimental design.

### NK cell isolation and culture

NK cells were isolated by negative selection from the spleens of naïve C57BL/6J mice using the EasySep Mouse NK Cell Isolation Kit (Stemcell Technologies, catalog no. 19855), according to the manufacturer’s instructions. Isolated NK cells were cultured in RPMI-1640 supplemented with 2.05mM L-glutamine, 10% FBS, 1% non-essential amino acids, 1% penicillin/streptomycin, 1% sodium pyruvate, 50 μM 2-beta-mercaptoethanol, 200ng/ml ng/mL recombinant mouse IL-2 and 20ng/ml ng/mL recombinant mouse IL-15 in a 5% CO2 cell culture incubator at 37°C for less than a week.

### NK cell cytotoxicity assay

The NK cell cytotoxicity assay was performed using the CytoTox96® Non-Radioactive Cytotoxicity Assay (Promega, catalog no. G1780), according to the manufacturer’s instructions. Target cells (5 × 10^3^) (primary YUMM1.7-GFP isolated from regressing tumors and residual disease, as mentioned above) in 50 μl of NK culture medium were seeded into each well of a 96-well non-treated cell culture plate. The effector cells (primary NK cells, as mentioned above) in 50 μl of NK culture medium were added at effector: target cell (E:T) ratios of 3:1 and 1:1, as indicated. Several controls were included, including Culture Medium Background Control, Effector Cell Spontaneous LDH Release Control, Target Cell Spontaneous LDH Release Control, Target Cell Maximum LDH Release Control, and Volume Correction Control. After four hours of incubation at 37°C, 50 μl of culture supernatants from all test and control wells were transferred to a fresh 96-well flat clear bottom plate to analyze LDH production. 50 μl of the CytoTox 96® Reagent was added to each well. After 30 minutes of incubation at room temperature, the absorbance at 492nm was recorded within 1 hour after adding the Stop Solution. After obtaining the corrected values by subtracting the absorbance value for the Culture Medium Background and Volume Correction Control, the percent cytotoxicity was calculated as follows:

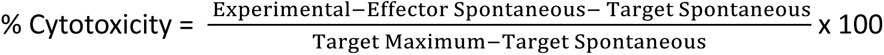

### Macrophage isolation and collection of macrophage-derived conditioned media

Macrophages were isolated by positive selection from regressing tumors or residual disease using the EasySep Mouse F4/80 Positive Selection Kit (Stemcell Technologies, catalog no. 100-0659) according to the manufacturer’s instructions. Isolated macrophages were cultured in complete macrophage medium (RPMI-1640 supplemented with 2.05mM L-glutamine, 10% FBS, 10% L-cell-conditioned medium, 1% penicillin/streptomycin, and 1% sodium pyruvate) at a seeding density of 2 x 10^6^ cells/3ml in complete macrophage medium in a 5% CO2 cell culture incubator at 37°C. After 22 hours, conditioned media was harvested and centrifuged at 1,500 x g for 10 minutes at 4°C to remove cell debris and stored at -20°C.

### Transwell assay

For NK cell migration in the context of L-1 treatment, NK cells were enriched from the spleens of naïve C57BL/6J mice, as described above. Enriched NK cells were treated with L-1 (1.5 μM) or an equivalent volume of DMSO as a control in NK culture medium (RPMI-1640 supplemented with 2.05mM L-glutamine, 1% non-essential amino acids, 1% penicillin/streptomycin, 1% sodium pyruvate, and 50 μM 2-beta-mercaptoethanol) with 0.5% FBS at 37°C and 5% CO2 for one hour. Treated NK cells were pipetted onto the insert of a 5μm pore size 24-well Transwell plate with a 6.5mm diameter insert (VWR, catalog no. 10769-236) at 2.5 × 10^5^ cells per 100 μl of medium in each well (two technical replicates). In the bottom chamber, 600 μl of NK culture medium with 10% FBS containing recombinant mouse Ccl5 (100 ng/ml) (Medchem Express, catalog no. HY-P71890) and Ccl8 (100 ng/ml) (Medchem Express, catalog no. HY-P7239) was added. The Transwell plate with treated NK cells was incubated at 37°C and 5% CO2 for four hours. After incubation, the inserts were removed. Migrated NK cells were counted using a hemocytometer.

For NK cell migration in the context of CCR2/5 inhibition, primary splenocytes from naïve C57BL/6J mice were treated with cenicriviroc (CVC, 10 μM) or an equivalent volume of DMSO as a control and pipetted onto the insert of a 5μm pore size 24-well Transwell plate with a 6.5mm diameter insert (VWR, catalog no. 10769-236) at 1 × 10^6^ cells per 100 μl of complete NK culture medium in each well. In the bottom chamber, 600 μl of macrophage conditioned media from regressing tumors or residual disease was added. The Transwell plate was incubated at 37°C and 5% CO2 for two hours. After incubation, the inserts were removed. The medium in the bottom chamber containing migrated splenocytes was collected. To calculate migrated NK cells, migrated splenocytes were stained with an anti-CD16/32 Fc receptor blocker (BioLegend, catalog no. 156603), LIVE/DEAD Fixable Violet Dead Cell Stain Kit (Thermo Fisher, catalog no. L34964), CD45-PerCp-Cy5.5 (BioLegend, catalog no. 103132), CD3-FITC (BioLegend, catalog no. 100204), and NK1.1-PE-Cy7 (Thermo Fisher, catalog no. 25-5941-81) antibodies. One hundred microliters of Precision Count Beads (BioLegend, catalog no. 424902) were added to calculate the absolute NK cell numbers.

### L-1 synthesis and characterization

Compound synthesis and characterization was performed as previously described *(54)* and also provided in the Supplementary Materials.

### L-1 treatment

Mice received L-1 treatment at a dose of 10 mg/kg. L-1 was dissolved in DMSO (10%; v/v) and subsequently diluted into Cremophor-EL (Fisher, catalog no. 61791-12-6) (5%; v/v) in sterile PBS to a final volume of 200 μl and administered to mice twice daily by intraperitoneal injection. Vehicle control contained 10% DMSO and 5% Cremophor-EL in sterile PBS. Mice bearing melanoma tumors were treated with L-1 or vehicle control for the duration of the study, starting from day 10 of BRAF/MEKi treatment.

### Statistical analysis

Graphs and statistical analyses were performed using GraphPad Prism 10 (RRID: SCR_002798) and R Studio (RRID: SCR_000432). Graphs show mean ± SEM or mean ± SD, as annotated in the corresponding figure legend. The number of tumors or mice per group used in each experiment is annotated in the corresponding figure legend as n. Two-tailed Student t-tests were used when two groups were compared with each other. A one-way analysis of variance (ANOVA) test with multiple comparisons was used to compare three or more groups. For Kaplan-Meier curves, a log-rank Mantel-Cox test was performed. Correlations were determined using simple linear regression. The statistical methods are shown in the figure legends. A P value of less than 0.05 was considered significant.

## SUPPLEMENTAL MATERIALS

Document S1. Figures S1-S11 and Table S1

Document S2. L-1 synthesis and characterization, related to Methods

## ACKNOWLEDGMENTS

The authors thank Dr. Anushka Dongre, Dr. Ayshwarya Subramanian, and Dr. Colleen Lau for providing feedback on the manuscript, Dr. Julie Sahler for expertise in flow cytometry, the Cornell Center for Animal Resources and Education (CARE) for research animal husbandry, and the Cornell Biotechnology Resource Center (BRC) Genomics Facility (RRID:SCR_021727), Bioinformatics Facility (RRID:SCR_021757), and Flow Cytometry Facility (RRID:SCR_021740) for their help with scRNA-seq, bulk mRNA-seq data and flow cytometry, with special thanks to Dr. Peter Schweitzer, Dr. Jen Grenier, and Dr. Lydia Tesfa for providing technical support.

## AUTHOR CONTRIBUTIONS

CHH conceived and conceptualized the study, designed and conducted experiments, analyzed and interpreted the data, and wrote the manuscript. KJL performed bioinformatics analysis of the scRNA-seq and bulk RNA-seq data. JC and LRD conducted experiments. JL synthesized and characterized L-1. DK analyzed the IF staining data. ZYZ provided funding for L-1 synthesis. ACW conceived and conceptualized the study, reviewed and edited the manuscript, and provided funding.

## DATA AND MATERIALS AVAILABILITY

All data are available in the main text or supplementary materials. Sequencing data will be available from the NCBI GEO repository upon publication.

