## Supplemental materials for "Innate Immune Remodeling Drives Therapy Resistance via Macrophage–NK Cell Crosstalk"

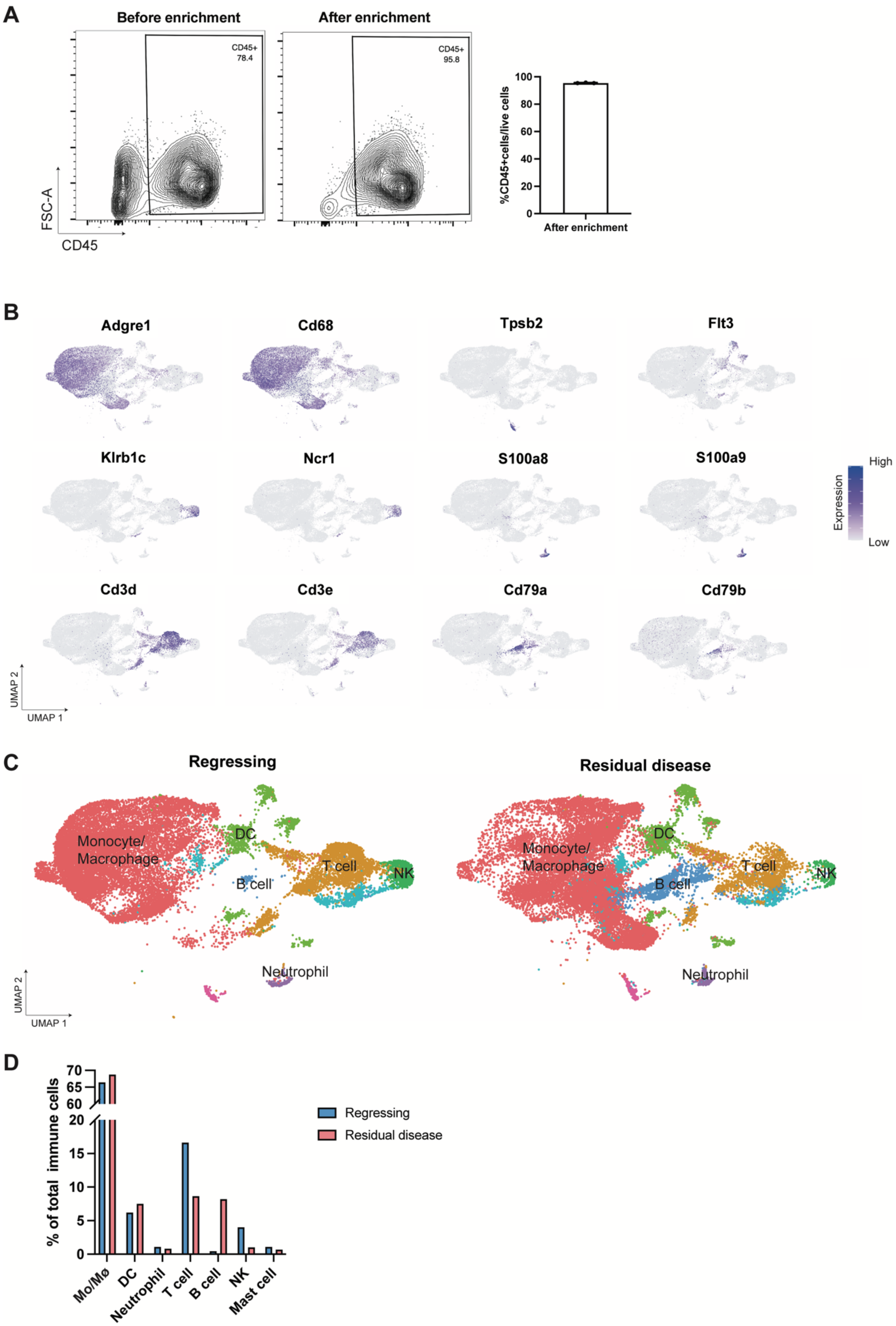
A
Before enrichment
After enrichment
CD45+
78.4
CD45+
95.8
FSC-A
CD45
%CD45+cells/live cells
After enrichment
B
Adgre1
Cd68
Tpsb2
Flt3
Klrb1c
Ncr1
S100a8
S100a9
Cd3d
Cd3e
Cd79a
Cd79b
High
Expression
Low
UMAP 2
UMAP 1
C
Regressing
Residual disease
Monocyte/
Macrophage
DC
T cell
B cell
NK
Neutrophil
D
% of total immune cells
Regressing
Residual disease
Mo/Mø
DC
Neutrophil
T cell
B cell
NK
Mast cell

**Figure S1. Immune cell types detected by single-cell RNA-seq in regressing tumors and residual disease.**

(A) Representative flow cytometry plot of CD45+ cells gated on live cells, before and after CD45 microbead enrichment. The numbers in the plots represent the percentage of cells within each gate. CD45+ cells represent nucleated hematopoietic cells. Bar graph showing the percentage of CD45+ cells out of live cells after CD45 microbead enrichment. N = 3 tumors.

(B) Expression of selected genes across the UMAP for immune cell-type annotation.

(C and D) UMAP plots showing immune cell types in regressing tumors and residual disease (C). Percentage of each immune cell type among total CD45+ cells in regressing tumors and residual disease (D).

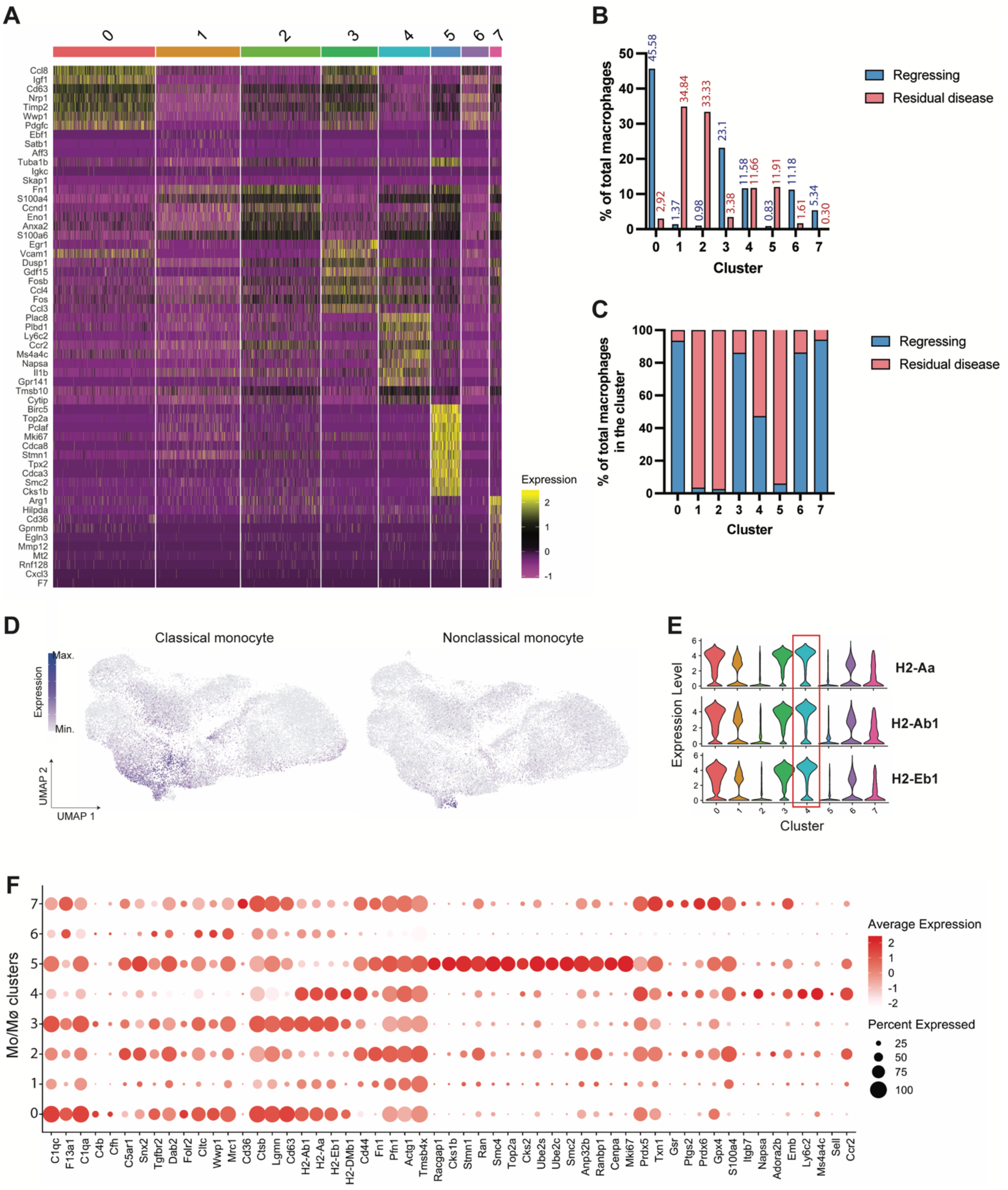


**Figure S2. Characterization of monocyte/macrophage populations detected by single-cell RNA-seq in regressing tumors and residual disease.**

(A) Heatmap of the top expressed genes within the eight monocyte/macrophage clusters.

(B and C) Percentage of monocytes/macrophages from regressing tumors and residual disease among total monocytes/macrophages (B) and within each cluster (C).

(D) Visualization of gene set scores associated with nonclassical/classical monocyte signatures.

(E) Expression of MHC II genes in each monocyte/macrophage cluster.

(F) Dot plot showing selected genes for complement & phagocytosis, antigen presentation, ECM & actin regulation, cycling, oxidative stress, and monocyte signatures in each macrophage cluster. Mo/Mϕ, monocyte/macrophage.

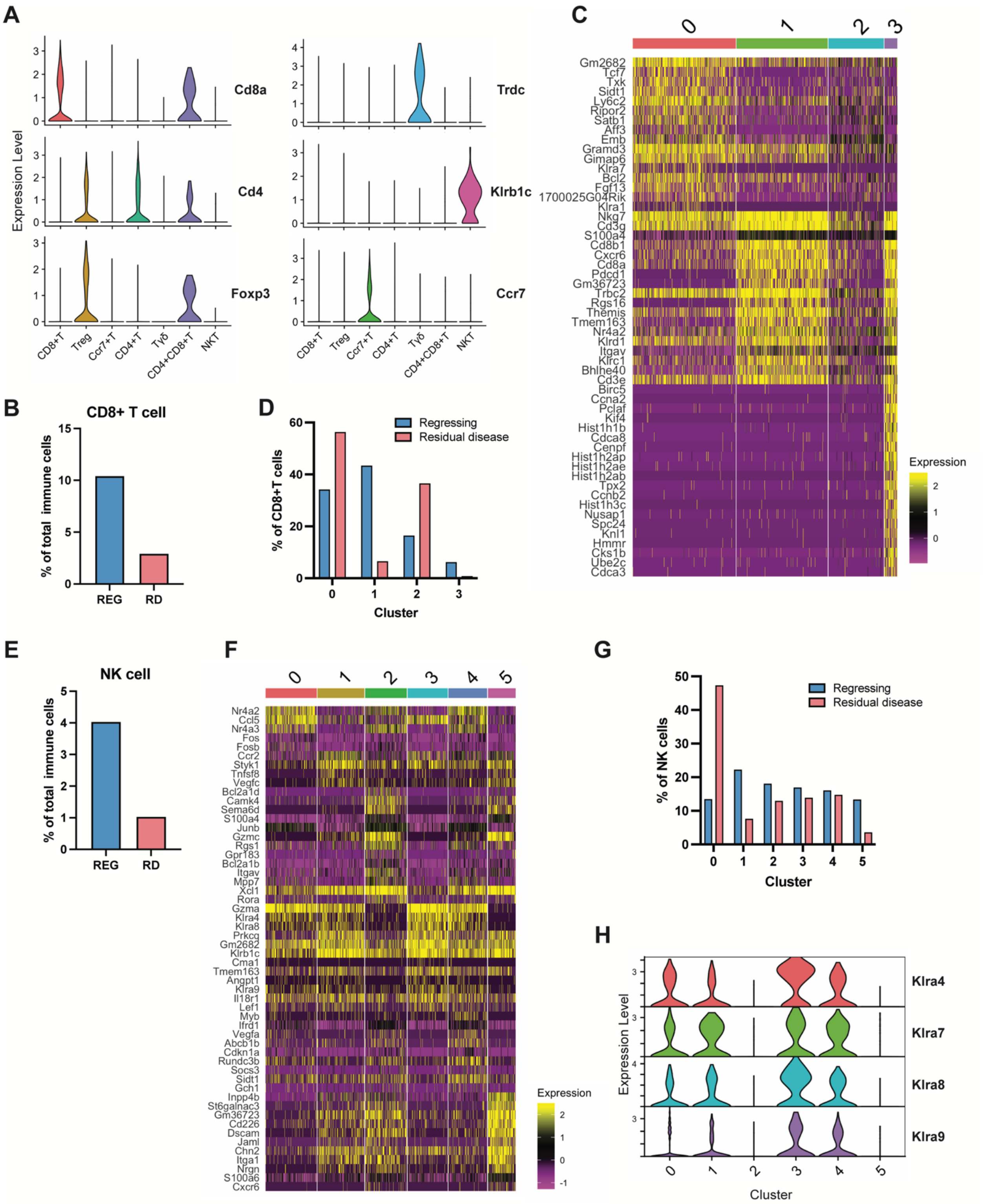


**Figure S3. Characterization of CD8+ T cell and NK cell populations detected by single-cell RNA-seq in regressing tumors and residual disease.**

(A) Expression of selected genes for T cell subtype annotation.

(B) Percentage of CD8+ T cells from regressing tumors and residual disease among total immune cells. REG, regressing; RD, residual disease.

(C) Heatmap of the top expressed genes within the four CD8+ T cell clusters.

(D) Percentage of CD8+ T cells from regressing tumors and residual disease among total CD8+ T cells.

(E) Percentage of NK cells from regressing tumors and residual disease among total immune cells. REG, regressing; RD, residual disease.

(F) Heatmap of the top expressed genes within the six NK cell clusters.

(G) Percentage of NK cells from regressing tumors and residual disease among total NK cells.

(H) Expression of Klra genes in each NK cell cluster.

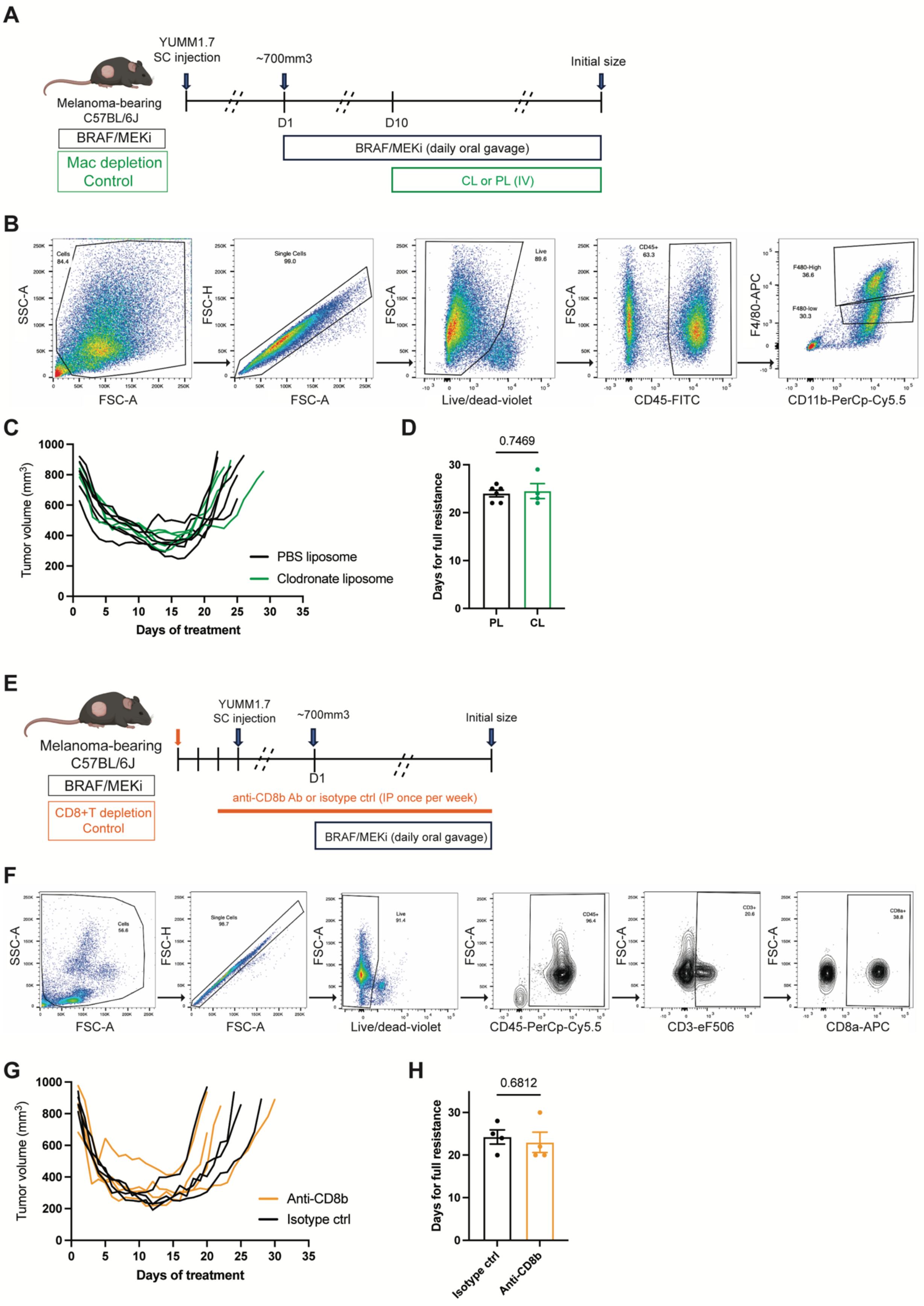

A
YUMM1.7
SC injection
~700mm3
Initial size
Melanoma-bearing
C57BL/6J
BRAF/MEKi
Mac depletion
Control
D1
D10
BRAF/MEKi (daily oral gavage)
CL or PL (IV)
B
SSC-A
FSC-A
Cells
84.4
FSC-H
Single Cells
99.0
Live
89.6
Live/dead-violet
CD45+
63.3
CD45-FITC
F4/80-APC
F480-High
36.6
F480-low
30.3
CD11b-PerCp-Cy5.5
C
Tumor volume (mm³)
Days of treatment
PBS liposome
Clodronate liposome
D
0.7469
Days for full resistance
PL
CL
E
YUMM1.7
SC injection
~700mm3
Initial size
Melanoma-bearing
C57BL/6J
BRAF/MEKi
CD8+T depletion
Control
D1
anti-CD8b Ab or isotype ctrl (IP once per week)
BRAF/MEKi (daily oral gavage)
F
SSC-A
FSC-A
Cells
56.6
FSC-H
Single Cells
98.7
Live
91.4
Live/dead-violet
CD45+
96.4
CD45-PerCp-Cy5.5
CD3+
20.6
CD3-eF506
CD8a+
38.8
CD8a-APC
G
Tumor volume (mm³)
Days of treatment
Anti-CD8b
Isotype ctrl
H
0.6812
Days for full resistance
Isotype ctrl
Anti-CD8b

**Figure S4. Macrophage depletion and CD8 + T cell depletion in BRAF/MEKi-treated mice to assess resistance onset.**

(A) Schematic illustrating macrophage depletion in BRAF/MEKi-treated mice bearing YUMM1.7. CL, clodronate liposomes; PL, PBS liposomes. Created with BioRender.com.

(B) Flow cytometry gating strategy for macrophages from BRAF/MEKi-treated tumors.

(C) Tumor volume curves of individual BRAF/MEKi-treated tumors following the administration of clodronate liposomes (CL) or PBS liposomes (PL).

(D) Days to resistance onset for BRAF/MEKi-treated tumors with administration of clodronate liposomes (CL) or PBS liposomes (PL). (n = 4-6 tumors, two-tailed unpaired t-test, mean $\pm$ SEM)

(E) Schematic illustrating CD8+ T cell depletion in BRAF/MEKi-treated mice bearing YUMM1.7. Created with BioRender.com.

(F) Flow cytometry gating strategy for CD8+ T cells from BRAF/MEKi-treated mice.

(G) Tumor volume curves of individual BRAF/MEKi-treated tumors after administering an anti-CD8b antibody or isotype control.

(H) Days to resistance onset for BRAF/MEKi-treated tumors with administration of an anti-CD8b antibody or isotype control. (n = 4 tumors, two-tailed unpaired t-test, mean $\pm$ SEM)

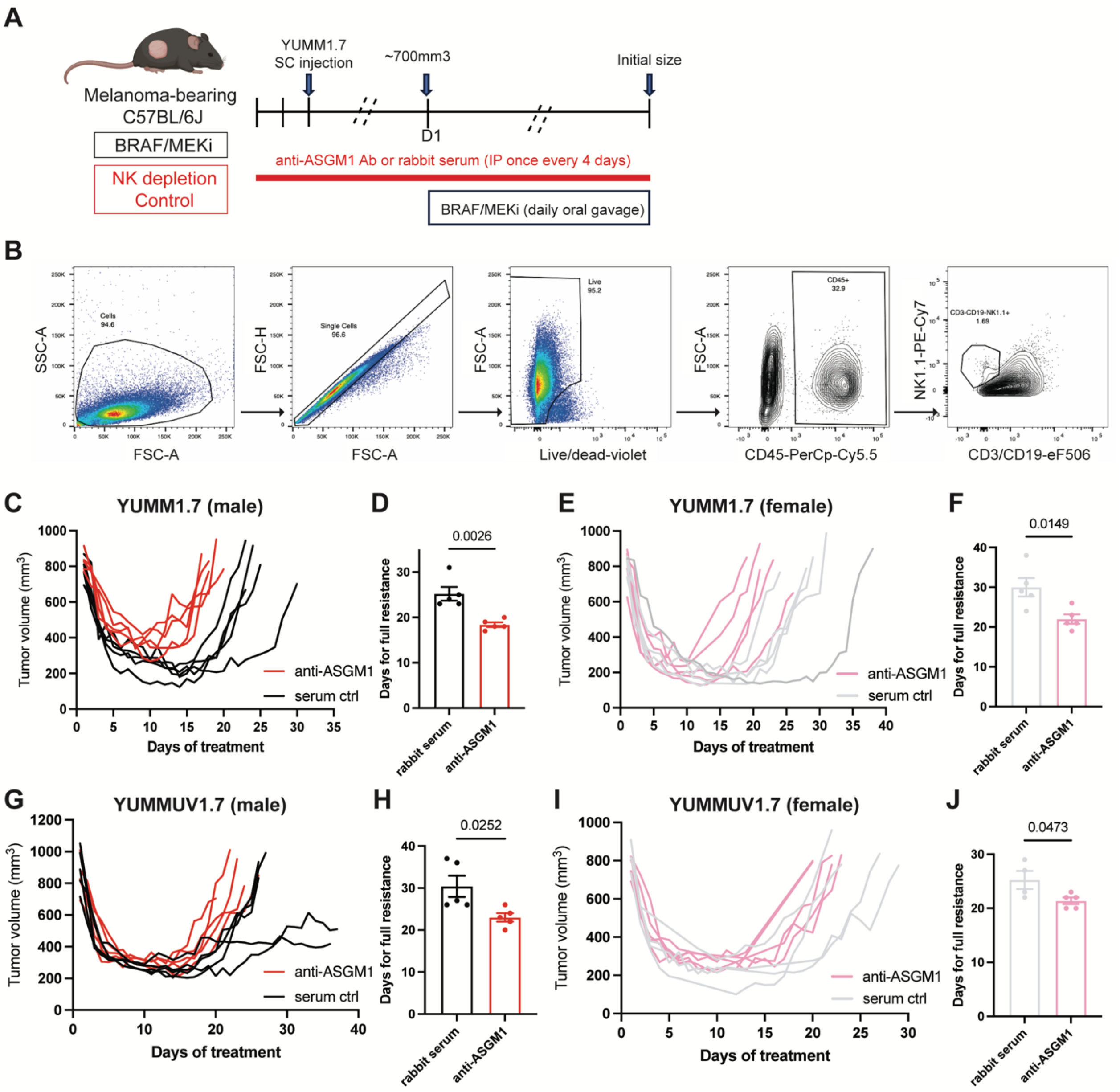


**Figure S5. NK cell depletion in BRAF/MEKi-treated mice to assess resistance onset.**

(A) Schematic illustrating NK cell depletion in BRAF/MEKi-treated mice bearing YUMM1.7. Created with BioRender.com.

(B) Flow cytometry gating strategy for NK cells from BRAF/MEKi-treated tumors.

(C) Tumor volume curves of individual BRAF/MEKi-treated YUMM1.7 in male mice with administration of an anti-ASGM1 antibody or rabbit serum.

(D) Days to resistance onset for BRAF/MEKi-treated YUMM1.7 in male mice with administration of an anti-ASGM1 antibody or rabbit serum. (n = 5 tumors, two-tailed unpaired t-test, mean ± SEM)

(E) Tumor volume curves of individual BRAF/MEKi-treated YUMM1.7 in female mice with administration of an anti-ASGM1 antibody or rabbit serum.

(F) Days to resistance onset for BRAF/MEKi-treated YUMM1.7 in female mice with administration of an anti-ASGM1 antibody or rabbit serum. (n = 5 tumors, two-tailed unpaired t-test, mean ± SEM)

(G) Tumor volume curves of individual BRAF/MEKi-treated YUMMUV1.7 in male mice with administration of an anti-ASGM1 antibody or rabbit serum.

(H) Days to resistance onset for BRAF/MEKi-treated YUMMUV1.7 in male mice with administration of an anti-ASGM1 antibody or rabbit serum to develop full resistance. (n = 5 tumors, two-tailed unpaired t-test, mean ± SEM)

(I) Tumor volume curves of individual BRAF/MEKi-treated YUMMUV1.7 in female mice with administration of an anti-ASGM1 antibody or rabbit serum.

(J) Days to resistance onset for BRAF/MEKi-treated YUMMUV1.7 in female mice with administration of an anti-ASGM1 antibody or rabbit serum. (n = 4-5 tumors, two-tailed unpaired t-test, mean ± SEM)

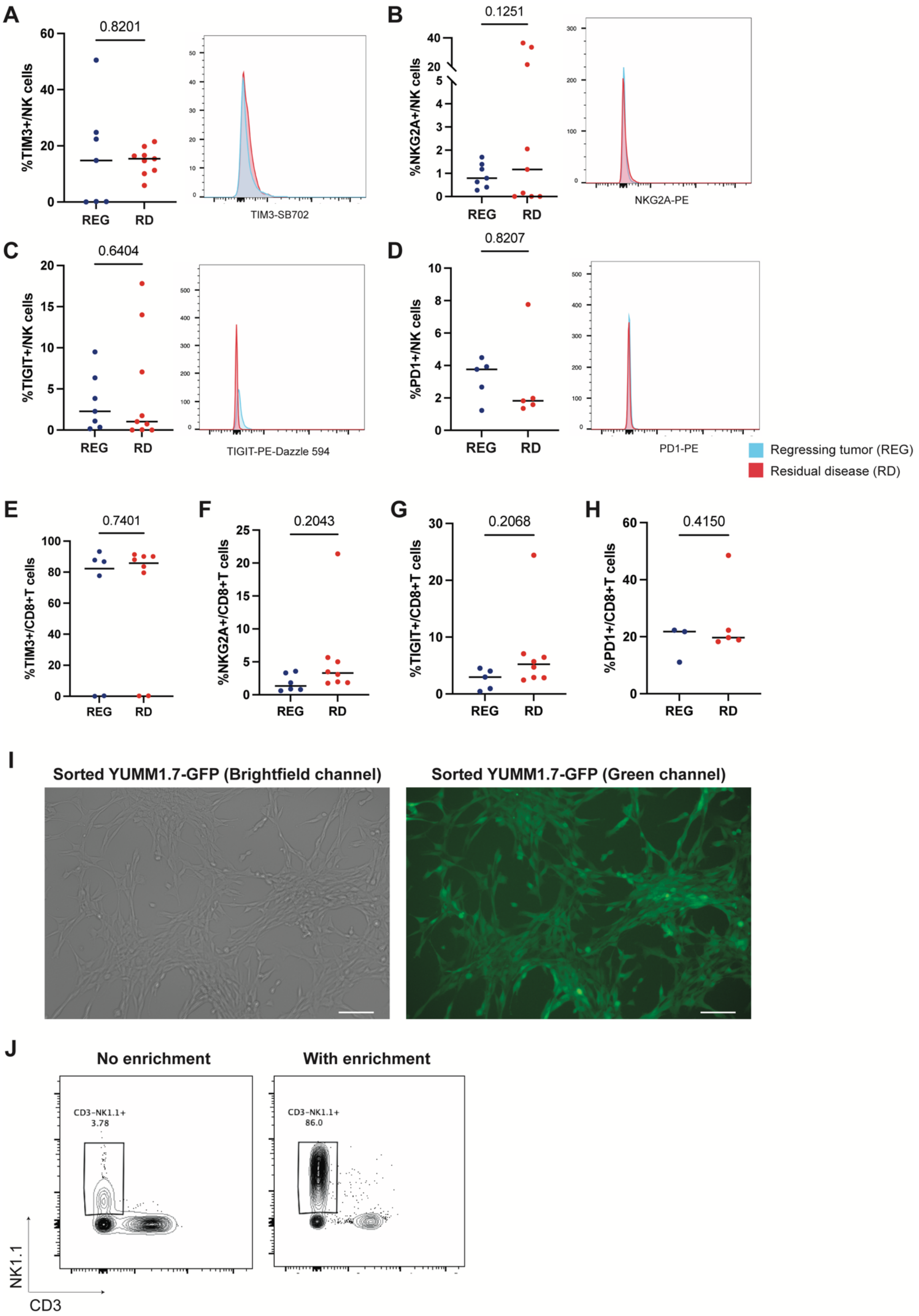

A
0.8201
%TIM3+/NK cells
REG
RD
TIM3-SB702
B
0.1251
%NKG2A+/NK cells
REG
RD
NKG2A-PE
C
0.6404
%TIGIT+/NK cells
REG
RD
TIGIT-PE-Dazzle 594
D
0.8207
%PD1+/NK cells
REG
RD
PD1-PE
Regressing tumor (REG)
Residual disease (RD)
E
0.7401
%TIM3+/CD8+T cells
F
0.2043
%NKG2A+/CD8+T cells
G
0.2068
%TIGIT+/CD8+T cells
H
0.4150
%PD1+/CD8+T cells
I
Sorted YUMM1.7-GFP (Brightfield channel)
Sorted YUMM1.7-GFP (Green channel)
J
No enrichment
With enrichment
CD3-NK1.1+
3.78
CD3-NK1.1+
86.0
NK1.1
CD3

**Figure S6. Exhaustion marker analysis in tumor-infiltrating NK cells, and enrichment for YUMM1.7-GFP and splenic NK cells.**

(A-D) Flow cytometry analysis of TIM3 (A), NKG2A (B), TIGIT (C), or PD1 (D) in tumor-infiltrating NK cells in regressing tumors and residual disease. REG, regressing; RD, residual disease. (n = 5-9 tumors, two-tailed unpaired t-test, median)

(E-H) Flow cytometry analysis of TIM3 (E), NKG2A (F), TIGIT (G), or PD1 (H) in tumor-infiltrating CD8+ T cells in regressing tumors and residual disease. REG, regressing; RD, residual disease. (n = 3-8 tumors, two-tailed unpaired t-test, median)

(I) Representative images of sorted YUMM1.7-GFP cells from BRAF/MEKi-treated tumors in culture.

(J) Representative flow cytometry plots for NK cells from spleen with or without enrichment, using negative selection.

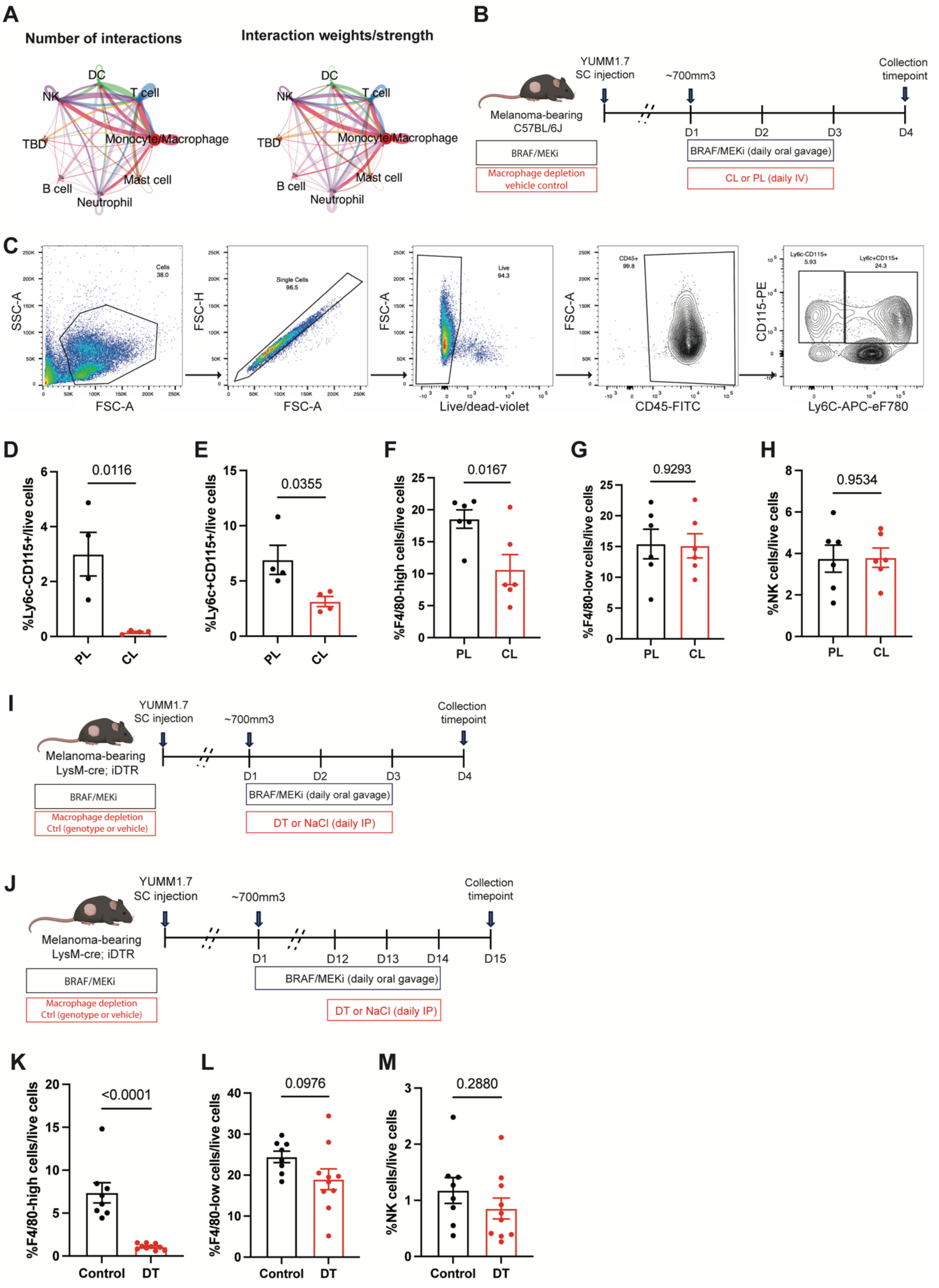

A
Number of interactions
Interaction weights/strength
DC
NK
T cell
TBD
Monocyte/Macrophage
Mast cell
B cell
Neutrophil
B
Melanoma-bearing
C57BL/6J
YUMM1.7
SC injection
~700mm3
Collection
timepoint
D1
D2
D3
D4
BRAF/MEKi
Macrophage depletion
vehicle control
BRAF/MEKi (daily oral gavage)
CL or PL (daily IV)
C
SSC-A
FSC-A
Cells
38.0
FSC-H
Single Cells
96.5
Live/dead-violet
Live
94.3
CD45-FITC
CD45+
99.8
CD115-PE
Ly6C-APC-eF780
Ly6c-CD115+
5.93
Ly6c+CD115+
24.3
D
%Ly6c-CD115+/live cells
0.0116
PL
CL
E
%Ly6c+CD115+/live cells
0.0355
F
%F4/80-high cells/live cells
0.0167
G
%F4/80-low cells/live cells
0.9293
H
%NK cells/live cells
0.9534
I
Melanoma-bearing
LysM-cre; iDTR
YUMM1.7
SC injection
~700mm3
Collection
timepoint
BRAF/MEKi
Macrophage depletion
Ctrl (genotype or vehicle)
BRAF/MEKi (daily oral gavage)
DT or NaCl (daily IP)
J
D1
D12
D13
D14
D15
K
%F4/80-high cells/live cells
<0.0001
Control
DT
L
%F4/80-low cells/live cells
0.0976
M
%NK cells/live cells
0.2880

**Figure S7. CellChat and macrophage analyses in tumor regression and residual disease.**

(A) Network diagrams depicting the communication interactions between each immune cell type during tumor regression. In the network diagram of Number of Interactions, the thickness of the lines indicates the number of interactions, with thicker lines likely representing more interactions (left). In the network diagram of Interaction Weights/Strength, the thickness of the lines corresponds to the strength of the interactions, with thicker lines representing stronger interactions.

(B) Schematic illustrating macrophage depletion by clodronate liposomes (CL) or PBS liposomes (PL) in BRAF/MEKi-treated mice bearing YUMM1.7 during tumor regression. Created with BioRender.com.

(C) Flow cytometry gating strategy for circulating monocytes from BRAF/MEKi-treated mice.

(D and E) Quantification of circulating Ly6c-CD115+ (D) or Ly6c+CD115+ (E) monocytes in BRAF/MEKi-treated mice receiving clodronate liposomes (CL) or PBS liposomes (PL) on day 4. (n = 4 mice, two-tailed unpaired t-test, mean ± SEM)

(F and G) Depletion efficiencies of clodronate liposomes in tumors during tumor regression. Quantification of F4/80-high (F) and F4/80-low macrophages (G). CL, clodronate liposomes; PL, PBS liposomes. (n = 6 tumors, two-tailed unpaired t-test, mean ± SEM)

(H) Quantification of tumor-infiltrating NK cells in BRAF/MEKi-treated mice receiving clodronate liposomes (CL) or PBS liposomes (PL) during tumor regression. (n = 6 tumors, two-tailed unpaired t-test, mean ± SEM)

(I and J) Schematic illustrating macrophage depletion in BRAF/MEKi-treated mice receiving diphtheria toxin (DT) (LysM-cre;iDTR) or control (NaCl in LysM-cre;iDTR or DT in WT mice) during tumor regression (I) or residual disease (J). Created with BioRender.com.

(K and L) Depletion efficiencies with diphtheria toxin (DT) (LysM-cre;iDTR) in tumors during residual disease. Quantification of F4/80-high macrophages (K) and F4/80-low macrophages (L). (n = 8-10 tumors, two-tailed unpaired t-test, mean ± SEM)

(M) Quantification of tumor-infiltrating NK cells in BRAF/MEKi-treated mice receiving diphtheria toxin (DT) (LysM-cre;iDTR) or control (NaCl in LysM-cre;iDTR or DT in WT mice) during residual disease. (n = 8-10 tumors, two-tailed unpaired t-test, mean ± SEM)

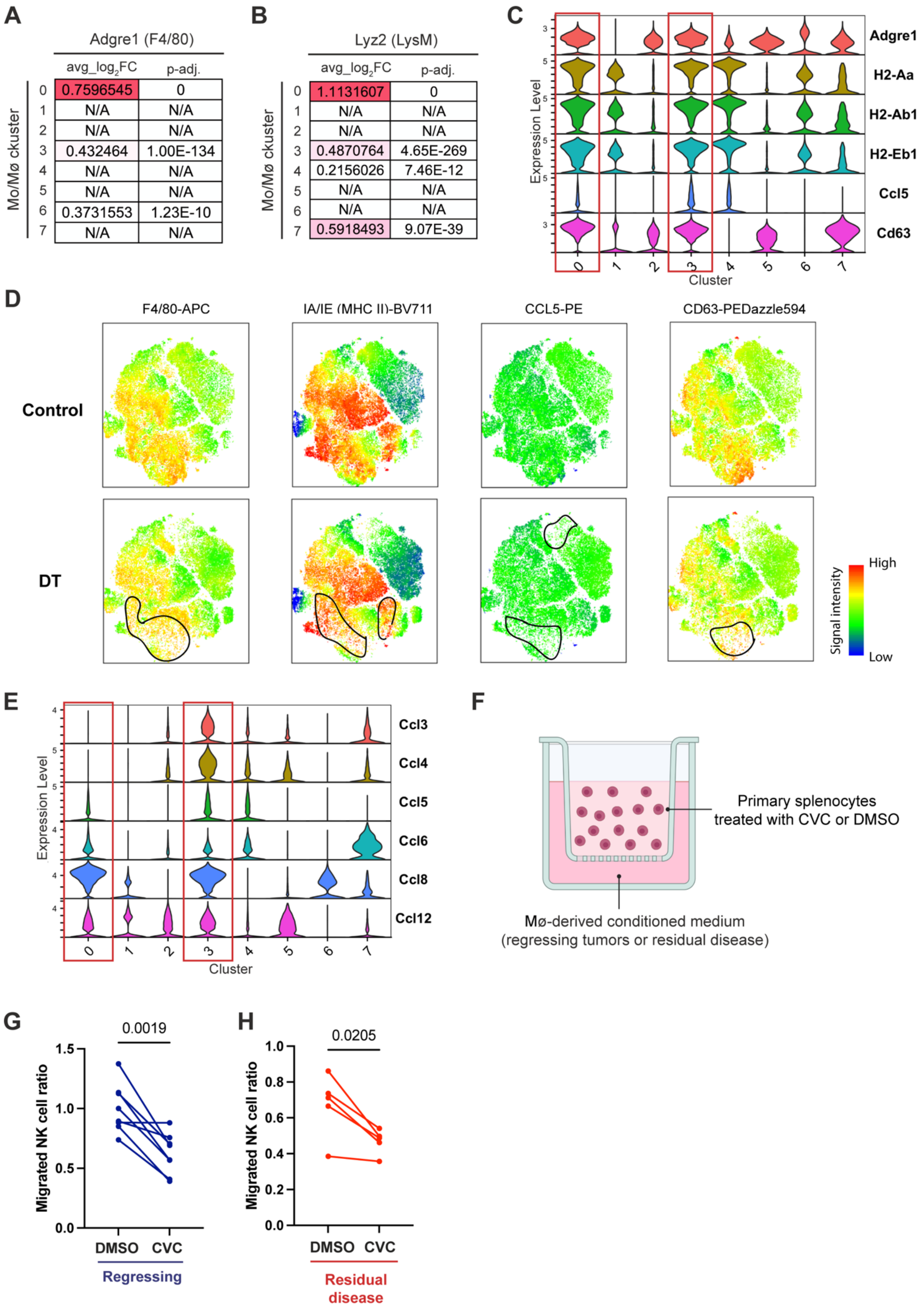
A
Adgre1 (F4/80)
avg_log2FC
p-adj.
Mo/Mø ckuster
0 0.7596545 0
1 N/A N/A
2 N/A N/A
3 0.432464 1.00E-134
4 N/A N/A
5 N/A N/A
6 0.3731553 1.23E-10
7 N/A N/A
B
Lyz2 (LysM)
avg_log2FC
p-adj.
Mo/Mø ckuster
0 1.1131607 0
1 N/A N/A
2 N/A N/A
3 0.4870764 4.65E-269
4 0.2156026 7.46E-12
5 N/A N/A
6 N/A N/A
7 0.5918493 9.07E-39
C
Expression Level
Adgre1
H2-Aa
H2-Ab1
H2-Eb1
Ccl5
Cd63
Cluster
D
F4/80-APC
IA/IE (MHC II)-BV711
CCL5-PE
CD63-PEDazzle594
Control
DT
Signal intensity
High
Low
E
Expression Level
Ccl3
Ccl4
Ccl5
Ccl6
Ccl8
Ccl12
Cluster
F
Primary splenocytes treated with CVC or DMSO
Mø-derived conditioned medium (regressing tumors or residual disease)
G
0.0019
Migrated NK cell ratio
DMSO
CVC
Regressing
H
0.0205
Migrated NK cell ratio
DMSO
CVC
Residual disease

**Figure S8. Analyses of specific macrophage populations and CCR2/5 axis in tumor regression.**

(A and B) Adgre1 (A) and Lyz2 (B) expression in each macrophage cluster. Mo/Mϕ, monocyte/macrophage.

(C) Expression of selected genes that upregulated in Clusters 0 and 3 in each macrophage cluster.

(D) Spectral flow cytometry on BRAF/MEKi-treated mice receiving diphtheria toxin (DT) (LysM-cre;iDTR) or control (NaCl in LysM-cre;iDTR) during tumor regression. tSNE of macrophage heterogeneity displaying selective markers that are upregulated in macrophage Clusters 0 and 3 in the scRNA-seq data.

(E) Expression of Ccl genes in each macrophage cluster.

(F) Schematic illustrating the experimental design of the transwell assay. CVC, cenicriviroc. Mϕ, macrophage.

(G and H) Paired dot plots showing the inhibiting effect of CVC on NK cell migration toward conditioned media derived from macrophages isolated from regressing tumors (G) and residual disease (H). (n = 5-8 tumors, conducted in two independent experiments, two-tailed paired t-test)

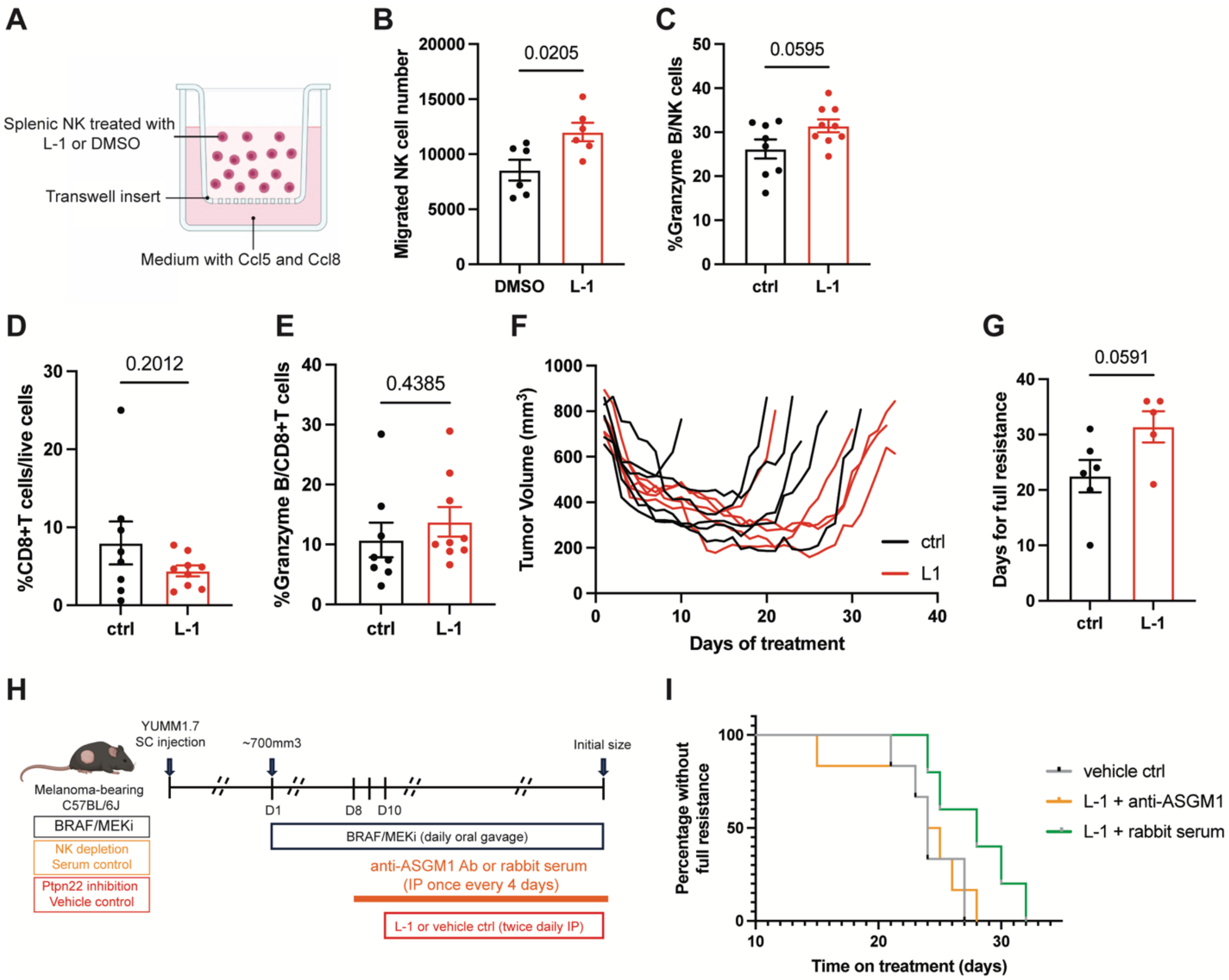


**Figure S9. Analysis of NK and CD8+ T cells and their correlation with the onset of resistance in mice treated with BRAF/MEK inhibitors and either L-1 or vehicle control.**

(A) Schematic illustrating the experimental design of the transwell assay. Created with BioRender.com.

(B) Quantification of migrated NK cells pretreated with L-1 or DMSO (vehicle control). (n = 3 biological samples, conducted in two independent experiments, two-tailed unpaired t-test, mean ± SEM)

(C) Quantification of Granzyme B in tumor-infiltrating NK cells from BRAF/MEKi-treated mice receiving L-1 or vehicle control on day 13. (n = 8-9 tumors, two-tailed unpaired t-test, mean ± SEM)

(D and E) Quantification of CD8+ T cells (D) or granzyme B in tumor-infiltrating CD8+ T cells (E) from BRAF/MEKi-treated mice receiving L-1 or vehicle control on day 13. (n = 8-9 tumors, two-tailed unpaired t-test, mean ± SEM)

(F) Tumor volume curves of individual BRAF/MEKi-treated tumors after administration of L-1 or vehicle control.

(G) Days to full resistance for BRAF/MEKi-treated tumors with L-1 or vehicle control administration. (n = 5-6 tumors, two-tailed unpaired t-test, mean ± SEM)

(H) Schematic illustrating the experimental design of NK cell depletion in YUMM1.7-bearing mice treated with BRAF/MEKi + L-1. Created with BioRender.com.

(I) Kaplan-Meier curve for BRAF/MEKi + L-1-treated mice bearing YUMM1.7 receiving an anti-ASGM1 antibody or rabbit serum. (n = 5-6 tumors)

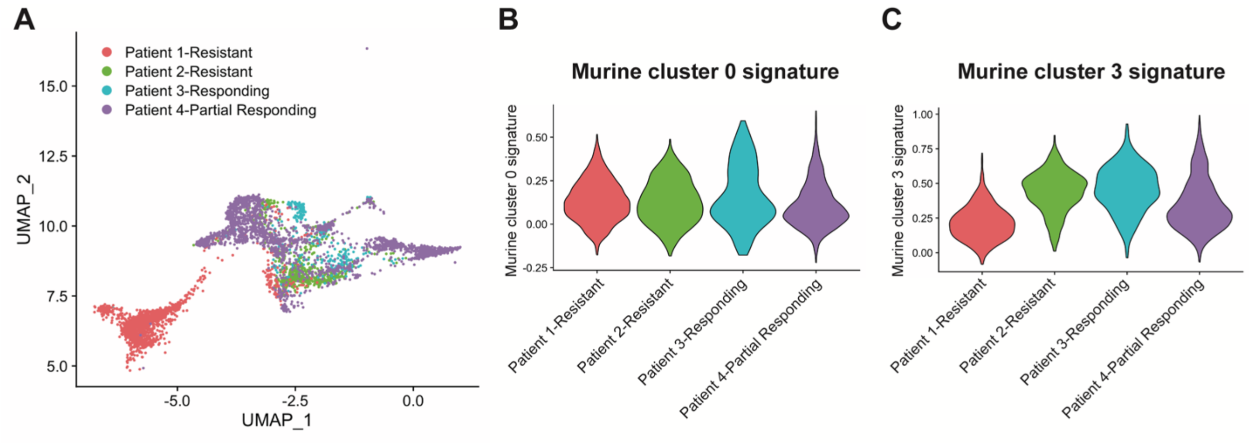


**Figure S10. Macrophage analysis of BRAF/MEKi-treated human melanoma fine-needle aspirate samples.**

(A) UMAP plot showing macrophage populations across the four patients.

(B and C) Violin plot showing murine macrophage Cluster 0 (B) or Cluster 3 (C) signature from Figure 1, displayed on four melanoma patient macrophage populations.

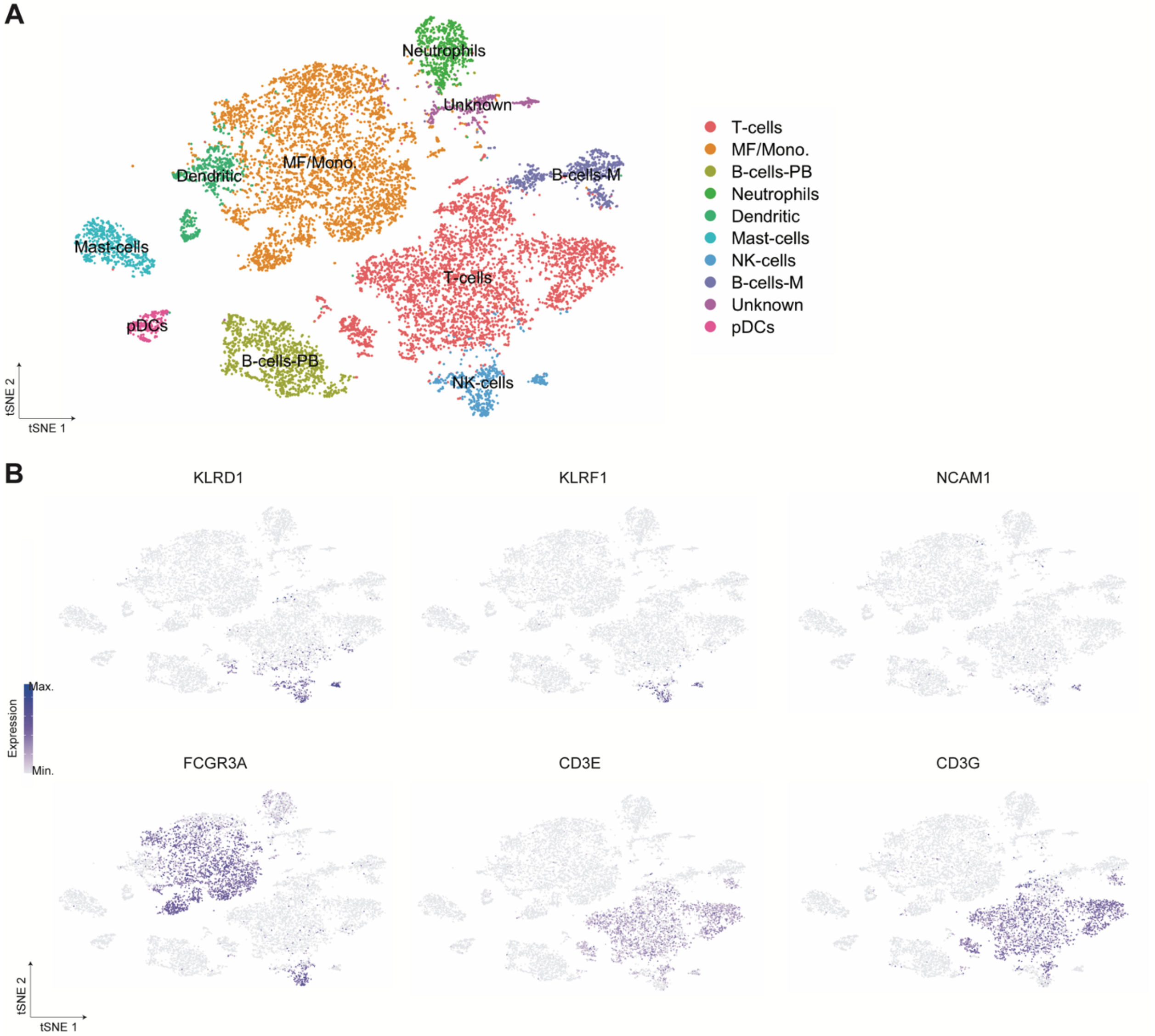


**Figure S11. Immune cell analysis of tyrosinase kinase inhibitor-treated human lung cancer biopsy samples.**

(A) tSNE plot showing immune cell populations across the all the samples.

(B) Expression of selected genes across the tSNE for NK cell annotation.

**Table S1. Antibodies**

| Antibodies | Source | Catalog Number |
|---|---|---|
| CD3 (17A2) | Biolegend | 100201 |
| NKR-P1C (NK1.1) (PK136) | Abcam | ab242134 |
| CD45-FITC (30-F11) | ThermoFisher Scientific | 11-0451-81 |
| CD45-PerCp-Cy5.5 (30-F11) | Biolegend | 103132 |
| CD3-FITC (17A2) | Biolegend | 100204 |
| CD3-eFlour 506 (17A2) | ThermoFisher Scientific | 69-0032-80 |
| CD19-eFlour 506 (EBIO1D3) | ThermoFisher Scientific | 69-0193-80 |
| CD8a-APC (53-6.7) | Biolegend | 100712 |
| CD115-PE | ThermoFisher Scientific | 12-1152-81 |
| CD11b-PerCp-Cy5.5 | ThermoFisher Scientific | 45-0112-80 |
| CD11b-FITC | Biolegend | 101205 |
| CCL5-PE (2E9/CCL5) | Biolegend | 149103 |
| F4/80-APC (BM8) | Biolegend | 123115 |
| Granzyme B- PE-Texas red (GB11) | ThermoFisher Scientific | GBR17 |
| I-A/I-E- Brilliant Violet 711 (M5/114.15.2) | Biolegend | 107643 |
| Ly6c-APC-efluor780 (HK1.4) | ThermoFisher Scientific | 47-5392-80 |
| NK1.1-PE-Cy7 (PK136) | ThermoFisher Scientific | 25-5941-81 |
| NKG2A$^{B6}$-PE (16A11) | Biolegend | 142803 |
| PD-1-PE (29F.1A12) | Biolegend | 135205 |
| S100B-PE (15F9NB) | Novus Biologicals | NBP2-45267PE |
| TIM-3-SB702 (RMT3-23) | ThermoFisher Scientific | 67-5870-82 |
| TruStain FcX Plus (anti-mouse cd16/32) (S17011E) | Biolegend | 156603 |

**General synthetic procedures and reagents.**

Unless otherwise specified, all reagents were purchased from commercial suppliers and used directly without further purification. Column chromatography was performed using KP-SIL silica gel (Biotage, USA), and flash column chromatography was performed on Biotage prepacked columns using the automated flash chromatography system Biotage Isolera One. The $^{1}H$ and $^{13}C$ NMR spectra were recorded on a Bruker AVANCE 500 MHz instrument. Chemical shifts for Proton magnetic resonance spectra ($^{1}H$ NMR) were quoted in parts per million (ppm) referenced to the appropriate solvent peak or 0.0 ppm for tetramethylsilane (TMS). The following abbreviations were used to describe peak splitting patterns when appropriate: br = broad, s = singlet, d = doublet, t = triplet, q = quartet, m = multriplet, dd = doublet of doublet. Coupling constants, J, were reported in hertz unit (Hz). Chemical shifts for 13C NMR were reported in ppm referenced to the center line at 39.52 of DMSO-d. Low-resolution mass spectra and purity data were obtained using an Agilent Technologies 6470 series, triple quadrupole LC/MS. High-resolution mass spectra (HRMS) were recorded on an Agilent Mass spectrometer using ESI-TOF (electrospray ionization-time of flight).

**Synthesis of Compound L-1**

Compound (1) served as the starting material and was condensed with Fmoc-L-Ala-OH using HOBt, HBTU, and DIPEA in DMF to yield the Fmoc-protected intermediate (2). The Fmoc group was subsequently removed with a 20% piperidine solution in DMF at room temperature. The resulting compound (3) was then condensed with biphenyl-4-carboxylic acid under similar reaction conditions as those used for compound (1), leading to the formation of compound (4). Finally, compound (4) was hydrolyzed using aqueous KOH in a methanol/water mixture and subsequently acidified with aqueous HCl to produce the final product, L-1, as the free acid.[1]

**Scheme 1.** Synthesis of Compound L-1[a]

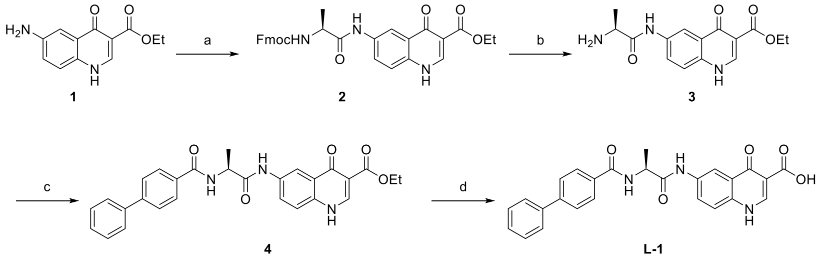


[a]Reagents and conditions: (a) Fmoc-L-Ala-OH, HOBt, HBTU, DIPEA, DMF, r.t., overnight, 68%; (b) Piperidine, DMF, r.t., 1 h, 70%; (c) Biphenyl-4-carboxylic acid, HOBt, HBTU, DIPEA, DMF, r.t., overnight, 75%; (d) KOH, MeOH/$H_2O$, 60°C, 16 h, 92%.

*(S)-ethyl 6-(2-((((9H-fluoren-9-yl)methoxy)carbonyl)amino)propanamido)-4-oxo-1,4-dihydroquinoline-3-carboxylate (**2**).*

Fmoc-L-Ala-OH (4.0 g, 12.84 mmol), HOBt (2.26 g, 16.7 mmol), and HBTU (6.34 g, 16.7 mmol) were dissolved in dry dimethylformamide (DMF, 80 mL). The mixture was stirred at room temperature for 15 minutes. Ethyl 6-amino-4-oxo-1,4-dihydroquinoline-3-carboxylate (2.68 g, 11.46 mmol) and N,N-Diisopropylethylamine (6.8 mL, 38.54 mmol) were then added, and the resulting mixture was stirred at room temperature overnight. The DMF was removed under reduced pressure using a rotary evaporator. Ethyl acetate and water were then added to the residue. The precipitate that formed was collected by filtration and purified by column chromatography, eluting with dichloromethane/methanol (10:1 v/v), to yield the Fmoc-protected intermediate (2) as a light brown solid (4.1 g, 68% yield). 1H NMR (500 MHz, DMSO) δ 12.28 (d, *J* = 6.4 Hz, 1H), 10.27 (s, 1H), 8.47 (d, *J* = 6.5 Hz, 1H), 8.39 (d, *J* = 2.1 Hz, 1H), 7.95 (dd, *J* = 8.9, 2.3 Hz, 1H), 7.88 (d, *J* = 7.6 Hz, 2H), 7.75 – 7.68 (m, 3H), 7.57 (d, *J* = 8.9 Hz, 1H), 7.44 – 7.37

(m, 2H), 7.35 – 7.28 (m, 2H), 4.28 – 4.26 (m, 2H), 4.21 – 4.17 (m, 4H), 1.32 (d, *J* = 7.1 Hz, 3H), 1.26 (t, *J* = 7.1 Hz, 3H). LC-MS (ESI): m/z $[M + H]^+$ calcd. For $C_{30}H_{28}N_3O_6$: 526.20, found: 526.30.

*(S)-ethyl 6-(2-aminopropanamido)-4-oxo-1,4-dihydroquinoline-3-carboxylate (**3**).*

The Fmoc-protected intermediate 2 (4.0 g, 7.61 mmol) was dissolved in DMF (60 mL). Piperidine (15.0 mL) was added, and the reaction mixture was stirred at room temperature for 1 hour. After concentration under vacuum, a brown solid was obtained. This solid was washed with ethyl acetate to afford the title compound (1.62 g, 70% yield). $^1$H NMR (500 MHz, DMSO) δ 8.49 (s, 1H), 8.43 (d, *J* = 2.4 Hz, 1H), 7.98 (dd, *J* = 8.9, 2.4 Hz, 1H), 7.58 (d, *J* = 8.9 Hz, 1H), 4.21 (q, *J* = 7.1 Hz, 2H), 3.52 – 3.47 (m, 1H), 1.27 (t, *J* = 7.1 Hz, 3H), 1.25 (d, *J* = 6.9 Hz, 3H). LC-MS (ESI): m/z $[M + H]^+$ calcd. For $C_{15}H_{18}N_3O_4$: 304.13, found: 304.20.

*(S)-ethyl 6-(2-([1,1'-biphenyl]-4-ylcarboxamido)propanamido)-4-oxo-1,4-dihydroquinoline-3-carboxylate (**4**).*

Biphenyl-4-carboxylic acid (0.80 g, 4.04 mmol), HOBt (0.7 g, 5.24 mmol), and HBTU (2.0 g, 5.24 mmol) were dissolved in dry dimethylformamide (DMF, 40 mL). The mixture was stirred at room temperature for 15 minutes. (S)-ethyl 6-(2-aminopropanamido)-4-oxo-1,4-dihydroquinoline-3-carboxylate (1.1 g, 3.64 mmol) and N,N-Diisopropylethylamine (2.14 mL, 12.1 mmol) were then added, and the resulting mixture was stirred at room temperature overnight. The DMF was removed using a rotary evaporator. Ethyl acetate and water were then added to the residue. The precipitate that formed was collected by filtration and washed with ethyl acetate to yield (S)-ethyl 6-(2-([1,1'-biphenyl]-4-ylcarboxamido)propanamido)-4-oxo-1,4-dihydroquinoline-3-carboxylate (1.32 g, 75% yield). $^1$H NMR (500 MHz, DMSO) δ 12.30 (d, *J* = 6.7 Hz, 1H), 10.36 (s, 1H), 8.74 (d, *J* = 7.0 Hz, 1H), 8.48 (d, *J* = 6.7 Hz, 1H), 8.41 (d, *J* = 2.3 Hz, 1H), 8.03 (d, *J* = 8.4 Hz, 2H), 8.00 (dd, *J* = 8.9, 2.4 Hz, 1H), 7.79 (d, *J* = 8.4 Hz, 2H), 7.75 (d, *J* = 7.2 Hz, 2H), 7.59 (d, *J* = 8.9 Hz, 1H), 7.50 (t, *J* = 7.6 Hz, 2H), 7.41 (t, *J* = 7.4 Hz, 1H), 4.67 – 4.61 (m, 1H), 4.21 (q, *J* = 7.1 Hz, 2H), 1.48 (d, *J* = 7.2 Hz, 3H), 1.28 (t, *J* = 7.1 Hz, 3H). LC-MS (ESI): m/z $[M + H]^+$ calcd. For $C_{28}H_{26}N_3O_5$: 484.19, found: 484.20.

*(S)-6-(2-([1,1'-biphenyl]-4-ylcarboxamido)propanamido)-4-oxo-1,4-dihydroquinoline-3-carboxylic acid (**L-1**).*

To a solution of compound 4 (1.0 g, 2.06 mmol) in methanol (40 mL) and water (40 mL), KOH (1.16 g, 20.68 mmol) was added. The mixture was stirred at 60°C for 16 hours. It was then cooled to 0°C and carefully acidified with 1N HCl until the pH reached ~1. The resulting precipitate was collected by filtration and purified by HPLC to yield the desired product L-1 as an off-white solid (0.85 g, 92% yield). $^1$H NMR (500 MHz, DMSO) δ 10.53 (s, 1H), 8.83 (d, *J* = 6.8 Hz, 1H), 8.78 (d, *J* = 6.8 Hz, 1H), 8.65 (d, *J* = 2.4 Hz, 1H), 8.10 (dd, *J* = 9.1, 2.4 Hz, 1H), 8.04 (d, *J* = 8.5 Hz, 2H), 7.84 – 7.77 (m, 3H), 7.76 – 7.70 (m, 2H), 7.50 (t, *J* = 7.6 Hz, 2H), 7.45 – 7.39 (m, 1H), 4.68 – 4.61 (m, 1H), 1.49 (d, *J* = 7.2 Hz, 3H). $^{13}$C NMR (126 MHz, DMSO) δ 177.99 (s), 172.03 (s), 166.55 (s), 166.08 (s), 144.05 (s), 142.89 (s), 139.15 (s), 137.34 (s), 135.40 (s), 132.67 (s), 129.04 (s), 128.30 (s), 128.08 (s), 126.88 (s), 126.41 (s), 126.19 (s), 124.95 (s), 120.41 (s), 113.21 (s), 107.07 (s), 50.13 (s), 17.61 (s). LC-MS (ESI): m/z $[M - H]^-$ calcd. For $C_{26}H_{20}N_3O_5$: 454.14, found: 454.30. HRMS (ESI-TOF): m/z $[M - H]^-$ calcd. For $C_{26}H_{20}N_3O_5$: 454.1403, found: 454.1413; Purity: >95% (UV, λ = 254 nm).
